# Multiple-site diversification of regulatory sequences enables inter-species operability of genetic devices

**DOI:** 10.1101/771782

**Authors:** Angeles Hueso-Gil, Ákos Nyerges, Csaba Pál, Belén Calles, Víctor de Lorenzo

## Abstract

The features of the light-responsive cyanobacterial CcaSR regulatory node that determine interoperability of this optogenetic device between *Escherichia coli* and *Pseudomonas putida* have been examined. For this, all structural parts (i.e. *ho1* and *pcyA* genes for synthesis of phycobilin, the *ccaS/ccaR* system from *Synechocystis* and its cognate downstream promoter) were maintained but their expression levels and stoichiometry diversified by [i] reassembling them together in a single broad host range, standardized vector and [ii] subjecting the non-coding regulatory sequences to multiple cycles of directed evolution with random genomic mutations (DIvERGE), a recombineering method that intensifies mutation rates within discrete DNA segments. Once passed to *P. putida*, various clones displayed a wide dynamic range, insignificant leakiness and excellent capacity in response to green light. Inspection of the evolutionary intermediates pinpointed translational control as the main bottleneck for interoperability and suggested a general approach for easing the exchange of genetic cargoes between different species i.e. optimization of relative expression levels and upturning of subcomplex stoichiometry.

## Introduction

Many naturally-occurring biosystems respond to the presence of light by activating or repressing distinct gene sets. Typically, light-reacting proteins and regulatory nodes are found in photosynthetic organisms, such as plants, *algae* and several bacterial groups like cyanobacteria and other marine microorganisms that use light to pump protons for obtaining an extra energy supply^*1*^. The control of bacterial behavior using light has become a most useful asset in synthetic biology, as it allows an instant and non-disruptive on/off switching of genetic constructs. Light as an inducer is cheap, precise, fast and clean, avoiding problems of chemical residues in the culture media when the circuit needs to be turned off. One useful source of parts for engineering light-responsive switches is the two-component CcaSR system of cyanobacterial *Synechocystis* species^*2*^. This green light-responsive setup controls expression of the phycobilisome linker protein CpcG2. When 520 nm light hits the membrane sensor protein CcaS bound to cofactor phycocianobilin (PCB), the protein autophosphorylates and transfers the phosphoryl group to the CcaR transcription factor. This in turn binds the so-called G box of *P*_*cpcG2*_ promoter and activates its transcription by σ^70^-RNA polymerase^*2*^. PCB can be easily produced in non-cyanobacterial hosts from the *heme* group through the action of enzymes Ho1 and PcyA. On this basis, the system was adapted, reshaped and implemented for controlling gene expression in *E. coli* upon exposure to the appropriate wavelength^*3*^. Since then, a collection of growingly improved light switches has been developed to the same end derived from the CcaSR node, using as a host both *E. coli* and *Bacillus subtilis*^*4-6*^. Unfortunately, while *E. coli* is a good chassis for prototyping of genetic devices, its applicability to biotechnological endeavors in industry and the environment is limited^*7, 8*^ and other hosts of engineered circuits need to be considered. Yet, as the parameters and transfer functions that operate the constructs most often change with the biological context, optogenetic tools optimized for *E. coli* do not perform well—or at all— when transferred to other bacteria.

For adapting existing light switches—and any other genetic devices—to new hosts, the correct balance of the functional components for *nesting* in the new regulatory and biochemical network of the new recipient is crucial. This requires the exploration of an ample solution space for finding the best combination of expression parameters of multiple constituents in a given host^*9*^. Along this line Schmidl et al^*5*^ adopted a combinatorial assembly of ribosomal binding sites (RBSs) and a shortened version of the promoter *P*_*cpcG2*_ (called *P*_*cpcG2-172*_) for improving the dynamic range of the CcaSR-based light switch in *E. coli*, which brought about a more balanced expression of the components. But things become more challenging when the issue is interoperability of the same device between two different, distant bacterial species. One appealing recipient of such light-responsive modules is the soil bacterium *Pseudomonas putida*, an emerging chassis for Synthetic Biology-based metabolic engineering because of its excellent pre-evolved properties of stress resistance and high capacity to host strong redox reactions^*10, 11*^. But if intra-species optimization is demanding enough, inter-species portability adds a further screw turn to the problem, as the new context alters the multiple parameters involved for a usable performance.

In this work, we have adopted a novel strategy for making the popular CcaSR-based light switch usable in *P. putida*. This workflow relies on keeping intact the structural parts of the device but letting its regulatory DNA (i.e. their promoters and RBS) to navigate through a very large solution space. This was enabled by DIvERGE (directed evolution with random genomic mutations^*12*^), a recombineering method that causes hypermutation *in vivo* of multiple, pre-set DNA segments of a target replicon. Following diversification and a two-step screening process, expression levels of each of the CcaSR system component (*ccaS, pcyA, ho1, ccaR, P*_*cpc*G2-172_)^*5*^ re-balanced themselves for an optimal nesting in *P. putida*. This outcome not only solved a practical problem, but it also showcased a general stratagem for handling interoperability of SynBio devices between otherwise distant hosts.

## Results and Discussion

### Gross implantation of the CcaS-CcaR system into *P. putida*

The CcaSR light-sensitive, two-component system was discovered in cyanobacteria *Synechocystis* species^*2*^ and later adapted and used to control gene expression in *E. coli* under illumination with an adequate wavelength^*4, 5*^. In these works, the CcaSR system components and enzymes for PCB production were expressed independently from constitutive promoters in isolated transcriptional units, with the exceptions of *ho1* and *pcyA*, which together formed a two-gene operon. These parts were carried in two different narrow host range plasmids: pSR43.6 harbouring the cassette *ccaS-ho1-pcyA* and pSR58.6 bearing the segment *ccaR-msfGFP* (Supplementary Table S5). These plasmids encode resistance to antibiotics and origins of replication suitable for manipulation in *E. coli* but not adequate for the destination microorganism, *P. putida*. For overcoming this problem, we moved the two relevant cassettes from their original plasmids to broad-host range vectors pSEVA631 and pSEVA241, respectively^*13*^. These approximately preserved the copy number of every functional DNA segment of the original system and thus maintained the dose of each component (Fig. 1A).

**Figure 1.**
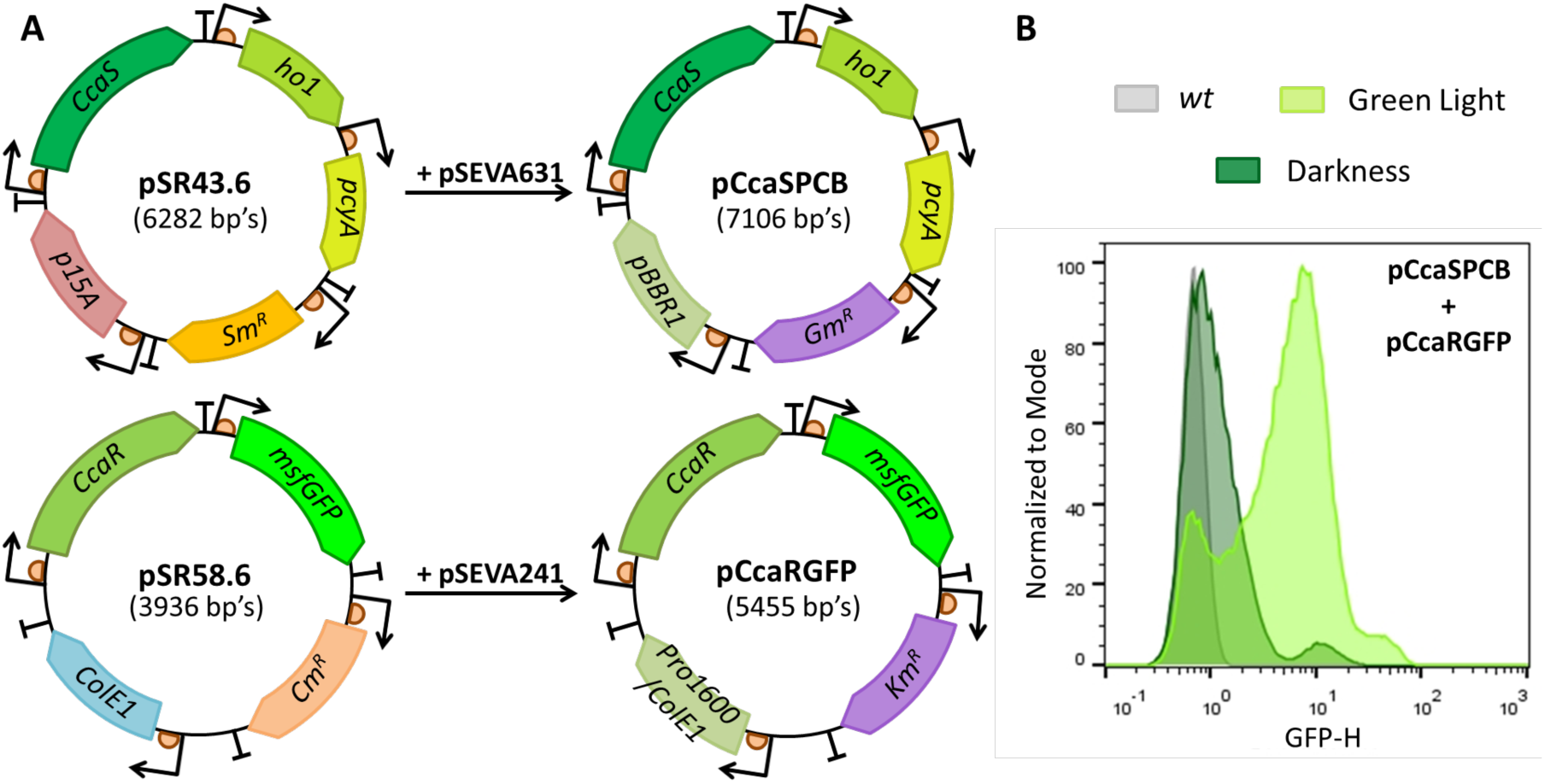
CcaSR two-component system original distribution and its behaviour. **A**. The functional DNA segments of the original vectors were adapted to the broad-host range pSEVA plasmids for their transfer to *P. putida*. Notice that the gene distribution was maintained and that the origins of replication of the destination backbone were selected to keep a similar plasmid copy number. **B**. Cell cytometry of *P. putida* bearing CcaSR-encoding plasmids after 8 h of induction. *P. putida* KT2440 *wt* with no GFP is plotted as a negative control (grey), together with *P. putida* harbouring pCcaSPCB + pCcaRGFP induced by green light (light green) and incubated in darkness (dark green).

The resulting constructs, pCcaSPCB (harbouring the DNA for *ccaS-ho1-pcyA*) and pCcaRGFP (cassette *ccaR-msfGFP*), were co-transformed into *P. putida* KT2440. This new strain was separately incubated under green light and under darkness conditions to check its functionality. After 8 h of incubation, the GFP fluorescent output was measured with flow cytometry, as this technology—unlike plate readersyields detailed information at single cell level. As shown in Fig. 1B, GFP expression indeed showed a response to the inducer (green light) when *P. putida* was transformed with both pCcaSPCB and pCcaRGFP plasmids. However, the mere adaptation of the antibiotic resistances and origins of replications to the new host was not enough to build an efficient system. Although basal levels were relatively low, fold induction was not as high as reported for *E. coli*^*5*^. Another interesting observation was a bimodal distribution of fluorescence reflected in two separate peaks both in darkness and in green light. In the non-induced state, there is a subpopulation displaying relative high fluorescence values while under inducing conditions a considerable amount of cells did not respond to illumination. In view of this clearly suboptimal performance we set out to achieve an homogeneous behavior, less leakiness, a higher dynamic range and capacity to better differentiate ON and OFF states of regulation.

In previous works with this system in *E. coli*, the key to obtaining a successful performance of the system involved balancing the production of each part of the optogenetic device^*5*^. For example, an excess of CcaR could provoke an unspecific phosphorylation of the regulator through other kinases and, therefore, unspecific transcription activation. On the other hand, low CcaR levels would not be enough for a differential GFP expression between light and darkness states. In addition, enzymes Ho1 and PcyA that produced the PCB cofactor could be poorly transcribed and/or translated. Moreover, there could be a bottleneck for the correct utilization of light by CcaS^*5*^. In any case, it was clear that due to the genetic context, parts and functional modules behaved unpredictably once interacting with the new cellular environment of *P. putida*. With these considerations in mind, we reasoned that it would be important to optimize the balance of CcaS/CasR for *P. putida* along with an adequate level of PCB biosynthesis.

In order to ease the tuning of the stoichiometry of the component of the device in *P. putida*, we first reassembled the whole set of parts (*ccaS, ho1, pcyA, ccaR* and *P*_*cpcG2-172*_ → *msfGFP*) in the same DNA segment and cloned it in two orientations in vector pSEVA621. Constructing two plasmids with opposite orientation was necessary because efficient diversification with DIvERGE (see below) requires the precise targeting of the replicating lagging strand with mutagenic oligonucleotides^*14, 15*^. As data on the structure of the replicore of RK2 origin-based expression vectors were not available, we overcome this issue by constructing and subsequently targeting two plasmids in opposing orientation. The resulting plasmids (Fig. 2A) carried such a cargo in either the same strand as the replication protein of RK2 origin, *trfA* (pGSFw) or in the other strand (pGSRv). As a consequence, both plasmids had the same functional segments, but oriented differently. The two constructs were then passed to *P. putida* and their responsiveness to green light tested.

**Figure 2.**
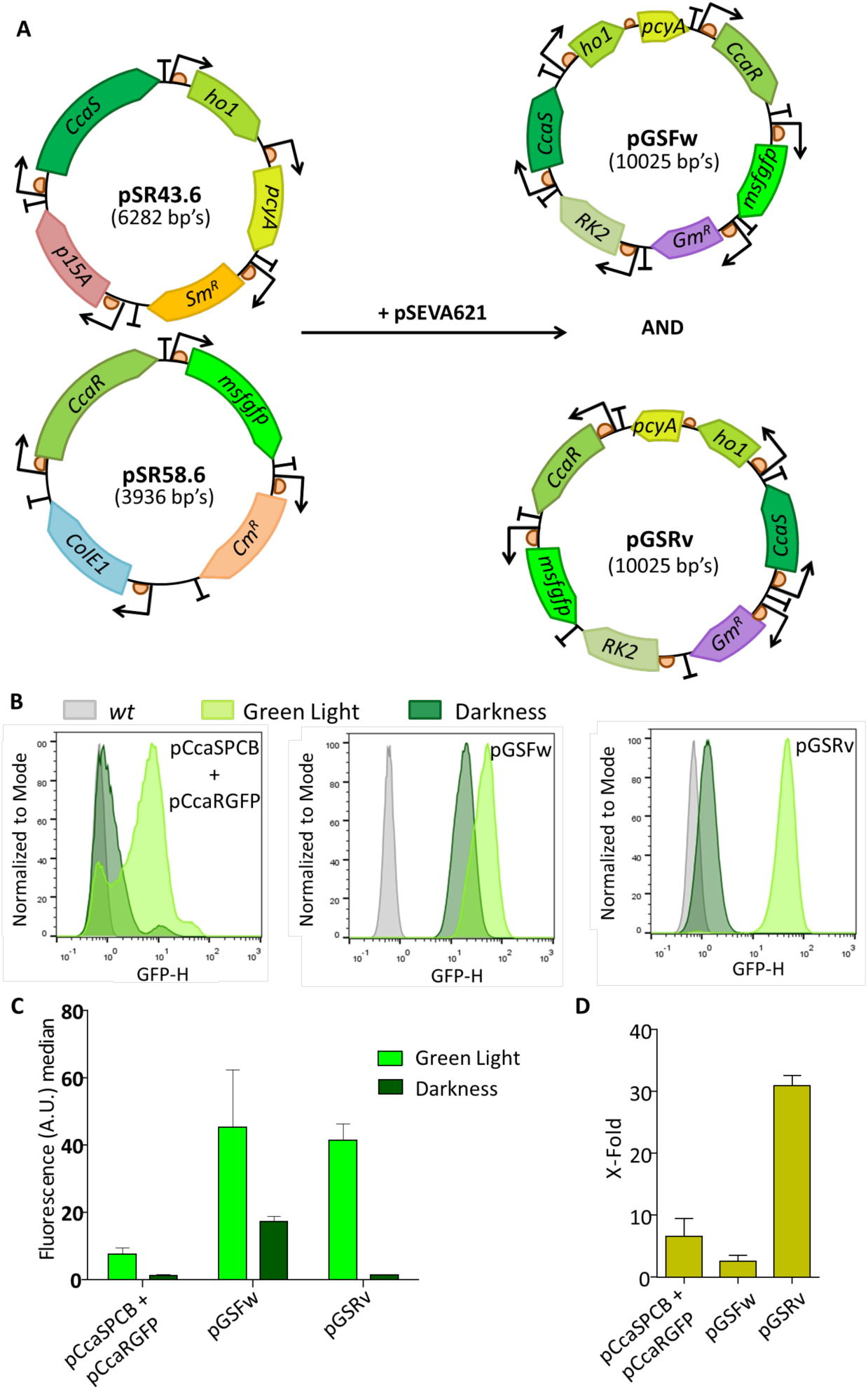
Behavior of a merged CcaSR-derived light-responsive device in *P. putida*. **A**. Second adaptation of the CcaSR system to *P. putida*.This time, both gene packs were merged in one single construct. The fused protein-coding cassette was placed in different orientations for the two final vectors: in the forward direction (pGSFw) and in its complementary counterpart (pGSRv). **B**. GFP expression profile of *P. putida* harbouring the CcaSR system in three plasmid formats. Cell cytometry for *P. putida* harbouring the two adapted plasmids, pCcaSPCB and pCcaRGFP, the single pGSFw plasmid with all the necessary genes in the forward orientation and the single pGSRv plasmid with same construct but in reverse orientation. In the three cases, fluorescence of *P. putida* KT2440 *wt* (grey curve), green light-induced cultures (light green) or darkness/non-induced cultures (dark green) are shown. **C**. Plotted medians for three flow cytometry biological replicates of light-induced and non-induced *P. putida* cultures with the same constructs. **D**. X-fold induction of the CcaSR system encoded in the three combinations of plasmids.

As shown in Fig. 2BC, cells bearing pGSFw and pGSRv could sense and respond to green light but they showed a very different behavior, both between them and in comparison with the 2-plasmid precursor system in which the relevant features of the device were split into two plasmids (Fig. 2B-D). Specifically, cells with pGSFw showed a high GFP production in the light-induced state, but also a high basal level without any illumination, meaning that its dynamic range and capacity was very low. In contrast, pGSRv showed a comparatively low basal level and a noticeable induction by light. These data exposed one more time the challenge of context dependency, specifically that the DNA strand carrying the coding information was making a considerable difference in the behavior of the thereby reassembled device. Although the ultimate basis of the dissimilarity between pGSFw and pGSRv remained speculative, the data of Fig. 2B-D suggested that the position of the first gene of our cassette, *ccaS*, could influence its transcription levels due to the upstream context, affecting the stoichiometry and behavior of the system. This advised to focus on the intergenic regulatory regions between genes as a way to optimize the behavior of the light switch in the new host.

### Diversification of regulatory regions of the CcaSR-based optogenetic device

Motivated by the results above, we set out to rebalance the expression level of each protein of the optogenetic device [*ccaS•pcyA/ho1•cca*R• *P*_*cpcG2-172*_ →] borne by pGSRv and pGSFw by introducing random mutations in their intergenic regulatory regions. Once such DNA segments were identified (*ccaS, pcyA/ho1, ccaR* and *P*_*cpcG2-175*_ RBSs andpromoters, see Fig. 3), we subjected them to directed evolution with random genomic mutations (DIvERGE; Nyerges *et al*., 2018), an ssDNA recombineering-based method that uses mutagenic oligonucleotides to integrate randomly distributed, random mutations to user-defined target regions, without inducing off-target mutagenesis. This approach involves the addition of mixtures of mutagenic oligonucleotides carrying randomly distributed point mutations that are incorporated into the target sequence upon invasion of the replication fork and integration as Okazaki fragments^*16, 17*^, facilitated by the β-recombinase of the Red system of phage λ^*18, 19*^. Such oligo-integration is easier in the lagging DNA strand which, as a result, has a higher mutation rate^*18, 20*^. In contrast, the leading strand has a more processive replication and integration of the oligonucleotides in the fork is less likely leading to a lower mutation rate. The primers used in our case were designed to target just one of the strands (Fig. 3A) and carried just 1 to 10 single point mutations per oligonucleotide molecule (i.e. they were soft-randomized^*12*^). Although we did not know upfront which one was the leading and which one was the lagging in the RK2 replicon of the SEVA plasmid, having pGSFw and pGSRv at hand covered both possibilities of hitting the right orientation of the DNA targets.

**Figure 3.**
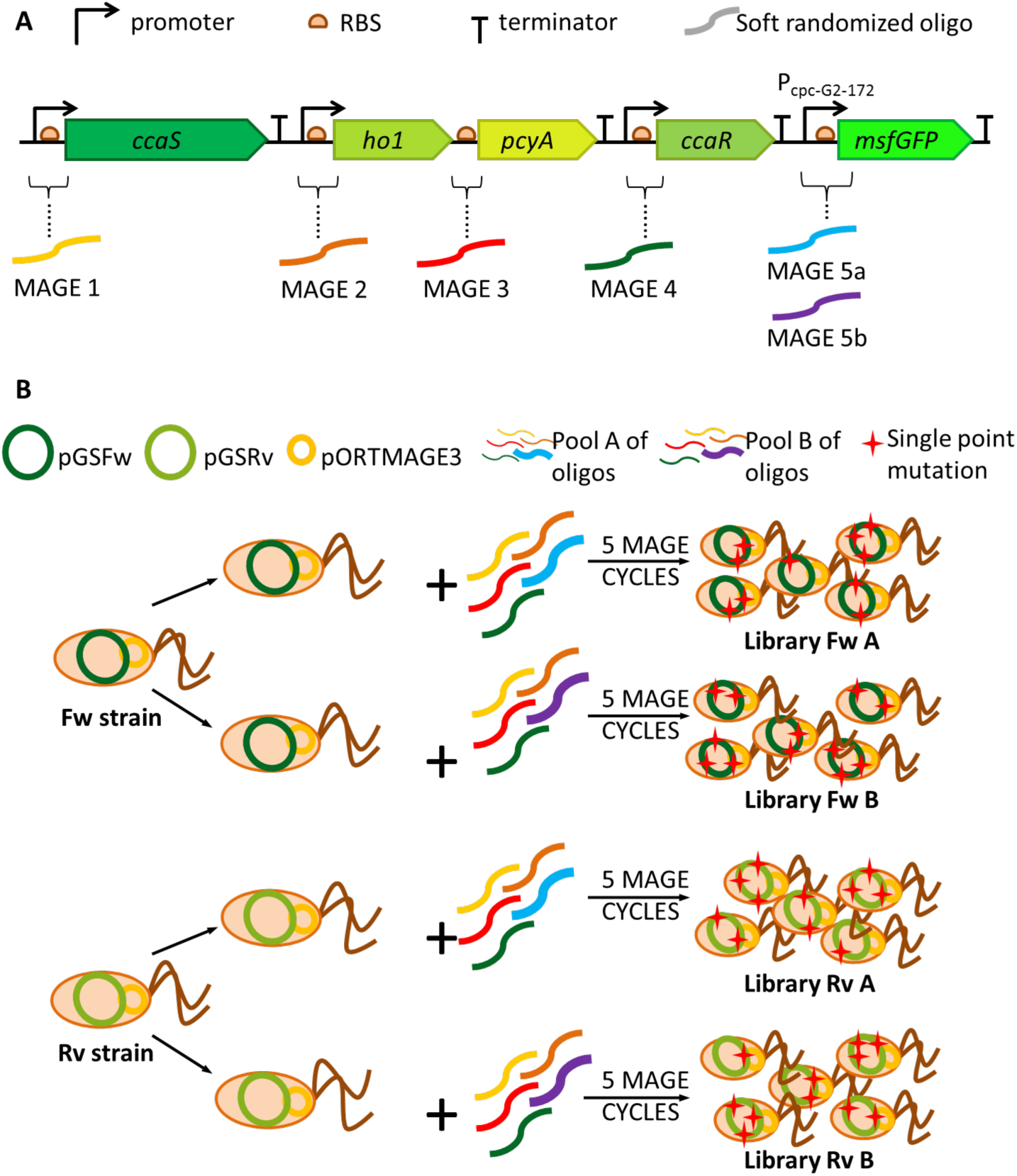
Strategy for diversification of regulatory sites. **A**. CcaSR cassette and DIVERGE oligo targets. Disposition of components as separate transcriptional units (except the two-gene operon *ho1-pcyA*) is shown. Each soft-randomized oligo mixture (including 1 or 2 mutations per molecule) was designed for the regulatory region of every gene including the promoters and the RBSs. For the p_cpc-G2-172_ sequence controlling the reporter gene, two different oligos were designed (DIVERGE 5a and DIVERGE 5b), targeting the G-Box recognized by CcaR or not including this sequence in the mutagenic process, respectively. **B**. Generation of CcaSR libraries using DIvERGE approach. Sketch of DIvERGE diversification over pGSFw and pGSRv plasmids using 2 pools of soft randomized oligos, carried out with *E. coli* MG1655 as host. The use of either mix of primers per each construct gave as a result 4 different libraries with random mutations in the regulatory regions of the CcaSR components. The four libraries are Fw A, Fw B, Rv A and Rv B.

Diversification of the targeted DNA sequences of pGSFw and pGSRv (Fig. 3) was carried out with *E. coli* MG1655 as a surrogate host for the procedure, bearing the pORTMAGE3^*19*^ plasmid with all the necessary machinery for the recombineering method. This includes a thermoinducible multicistronic operon encoding the Redβ recombinase and the negative dominant E32K variant of MutL that transiently suppresses the mismatch repair system and thus enables unbiased introduction of all possible nucleotide replacements^*21*^.

The result of 5 cycles of DIvERGE were two plasmid libraries per construct (A and B), each of which stemming from a different combination of mutagenic oligonucleotides. The dissimilarity between libraries A and B relies on oligo DIvERGE5, as it came in two versions. DIvERGE5a did not target the G-Box of the *P*_*cpcG2-172*_ promoter while DIvERGE5b did. The G-Box is a sequence motif (CACGTG) present in a large number of plant and cyanobacterial promoters that are recognized by a cognate transcription factor for expression control. In the device assembled in pGSFw and pGSRv this motif is bound by CcaR, what is followed by the recruitment of σ^70^-RNAP polymerase^*2, 22*^. By using two variants of DIvERGE5 we could either spare the G-box or try to improve it. On these basis, the target sequences of pGSFw plasmid were diversified using pool A of ssDNA oligonucleotides (DIvERGE1, DIvERGE2, DIvERGE3, DIvERGE4 and DIvERGE 5a), which resulted in the library Fw-A. The same plasmid subjected to pool B of oligos (DIvERGE1, DIvERGE2, DIvERGE3, DIvERGE4 and DIvERGE5b) produced library Fw-B (Fig. 3). By the same token, treatment of cells bearing pGSRv with oligo mixtures A or B generated library Rv-A and Rv-B, respectively (Fig. 3). Mutagenesis thus simultaneously reached out each promoter and RBS of every gene of the CcaSR-based system. Note that—as mentioned above—DIvERGE oligos were soft-randomized such that every ssDNA molecule was calculated to bear 1 to 25 indiscriminately distributed mutations compared to the parental, wild-type sequence (see Supplementary Table S7). This ensured efficient oligo-integration at the replication fork during the course of DIvERGE procedure. Based on the number of targeted cells and the average efficiency of pORTMAGE-based oligo integration (i.e., 30%), we have estimated that the effective library size approached approximately10^9^ variants per each library^*12*^. At this point, the population of *E. coli* bearing libraries Fw-A, Fw-B, Rv-A and Rv-B were separately pooled, lysed to extract the plasmid contents and the cccDNA fraction transferred to *P. putida* by transformation (see Materials and Methods) for securing a minimal loss of variability. The mere visual inspection of the resulting *P. putida* transformants (Supplementary Fig. S5) suggested that Rv libraries had a higher variability in their response to illumination, in contrast with the more homogeneous reaction to green light of the clones with the Fw. We interpreted this as an indication that the lagging strand of the RK2 origin of replication has the orientation of the antisense strand encoding the *repA* protein. It thus seemed that diversification was more efficient in the libraries generated with pGSRv plasmid, something surely related to the location of the lagging strand in vector pSEVA621.

### Isolation and enrichment of improved light-responsive devices in *P. putida*

In order to identify improved variants of the [*ccaS•pcyA/ho1•cca*R• *P*_*cpcG2-172*_ →] system with a lower basal expression level and a superior output in *P. putida*, we started by sorting the 4 libraries to isolate clones with the highest fluorescence levels in response to light induction. The sorting process involved three positive rounds where the best GFP emitters were selected after induction. This was then followed by a fourth negative cycle to collect clones with lower fluorescent emissions under darkness conditions, as detailed in the Material and Methods section. Total events selected per round of sorting and their relative percentages of the total events per population are shown in Supplementary Table S3. During the sorting process, we noticed that the 4 libraries evolved differently through the enrichment steps. As shown in Figure 4A, none of the Fw libraries (whether A or B versions) failed to display any meaningful shift in their cell cytometry profiles because of an improvement of their fluorescence. In contrast both Rv-A and Rv-B libraries were enriched in high-GFP variants after every sorting cycle. As a consequence, we chose only clones bearing Rv libraries for the rest of the screening, as the probability to find an improved device was higher because of its greater variability. On this basis, we isolated 2610 clones (1305 from the Rv-A library and 1305 from the Rv-B library) in 96-well plates with the help of a colony picker. By comparing with samples before sorting (Fig. 4A), we noticed that the enrichment process had significantly increased the number of clones that responded differentially to the presence of light (Fig. 4A, Supplementary Fig. S5). After GFP measurements under light induction and in darkness conditions of the 30 plates with the clones, we picked what appeared to be the best 84 clones. The two main parameters of these (basal level fluorescence and fold induction) were separately measured again in three biological replicates and plotted (Fig. 4B) for selecting the 5 best performers for further characterization.

**Figure 4.**
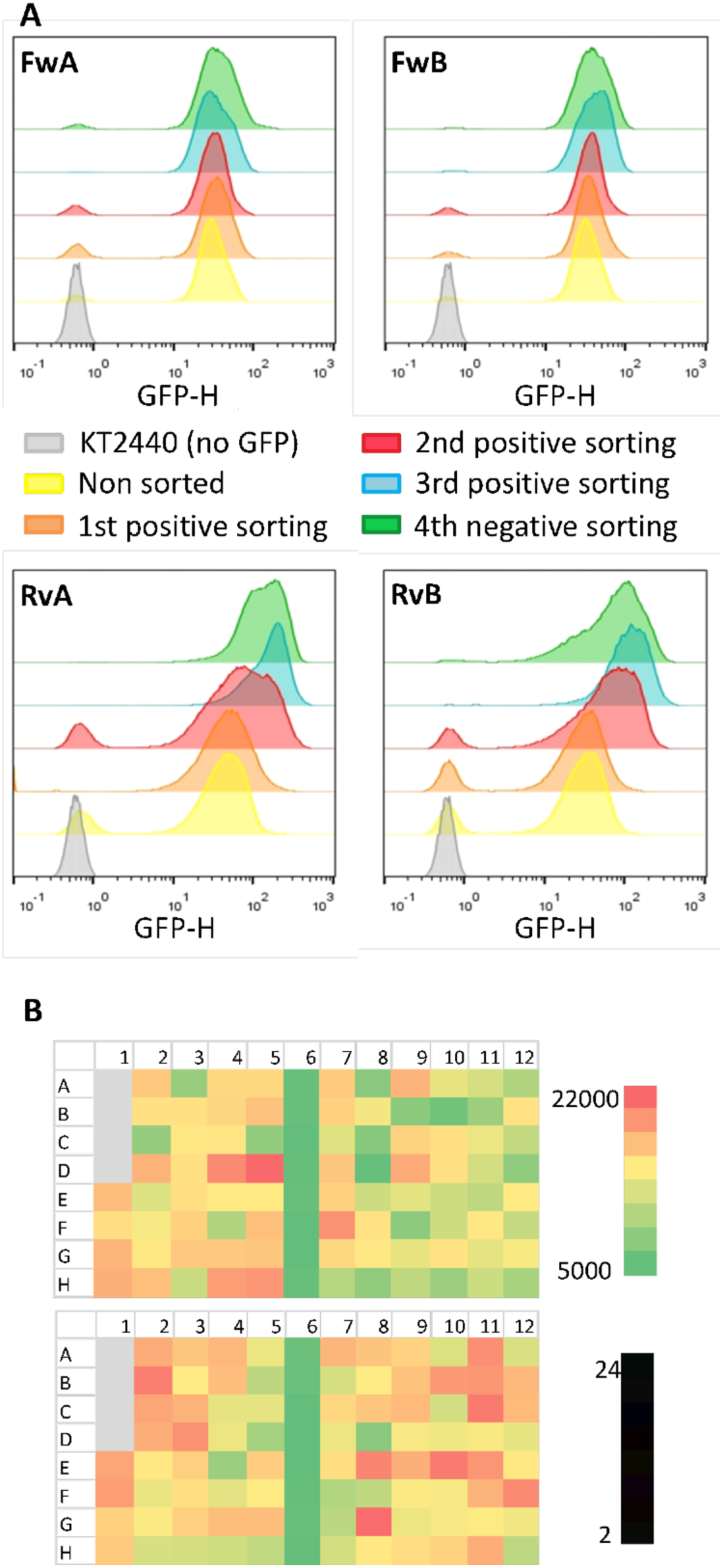
Analysis of libraries FwA, FwB, RvA and RvB. **A**. Population fluorescence of the four libraries before being sorted using FACS (yellow curve), after three sortings for selection of the induced population (first cycle, orange curve; second cycle, red curve; third cycle, blue curve) and after a fourth sorting event selecting the most GFP-negative non-induced population (green curve). **B**. Heatmaps for the final selected 84 clones representing the two criteria for selecting the best performers. On top, the basal values of fluorescence were plotted. The lower values are represented in green, while higher values correspond to red squares. In the bottom, X-fold induction values between the non-induced state and the light-induced state are shown. Higher X-fold values are in green while lower counterparts are plotted in red. In either heatmap, grey squares correspond to blank measurements, while column 6 was inoculated with the pGSRv-containing strain, thereby showing a uniform colour in both cases. Represented values are the calculated mean of three biological replicates. To select the 5 best clones, we chose wells with lower basal levels combined with higher X-fold induction.

### Phenotypic characterization of evolved light-responsive devices

The 5 clones mentioned above were taken through a suite of Flow Cytometry experiments. This technology not only allows the monitoring of ON/OFF states but also exposes homogeneity of gene expression throughout the population. The plasmids carried in those clones were named after the well where they were found in the cognate 96-well plate: pB10, pC8, pA8, pB11 and pB9 (Figure 4B). Comparing the results of the five clones among them and with those bearing the non-mutated construct of pGSRv, we noticed that pB9 originated the lowest fluorescence when cultures were not induced and had the widest dynamic range when exposed to green light. In order to confirm that these results were only due to the relevant parts of pB9, the DNA segment containing the complete evolved CcaSR system was reassembled again in the same vector backbone, pSEVA621, thereby yielding plasmid pGreenL (Fig. 5A-C), which had the same behavior as the parental one. *P. putida* cells carrying pGreenL were then subjected to further analyses. First, we inspected whether the system was specifically responding to green light and not to illumination by any other wavelength. In particular, we wished to check whether red light inhibited activity of the CcaSR system at levels below those in darkness, non-induction levels, as had been previously reported^*2, 5*^. To answer this question, we illuminated cultures of *P. putida* (pGreenL) with red light for 8 hours. The results shown in Supplementary Fig. 6B indicated that [i] the CcaSR sensor was blind to red light and [ii] the same red light had no effect on basal expression levels of GFP.

**Figure 5:**
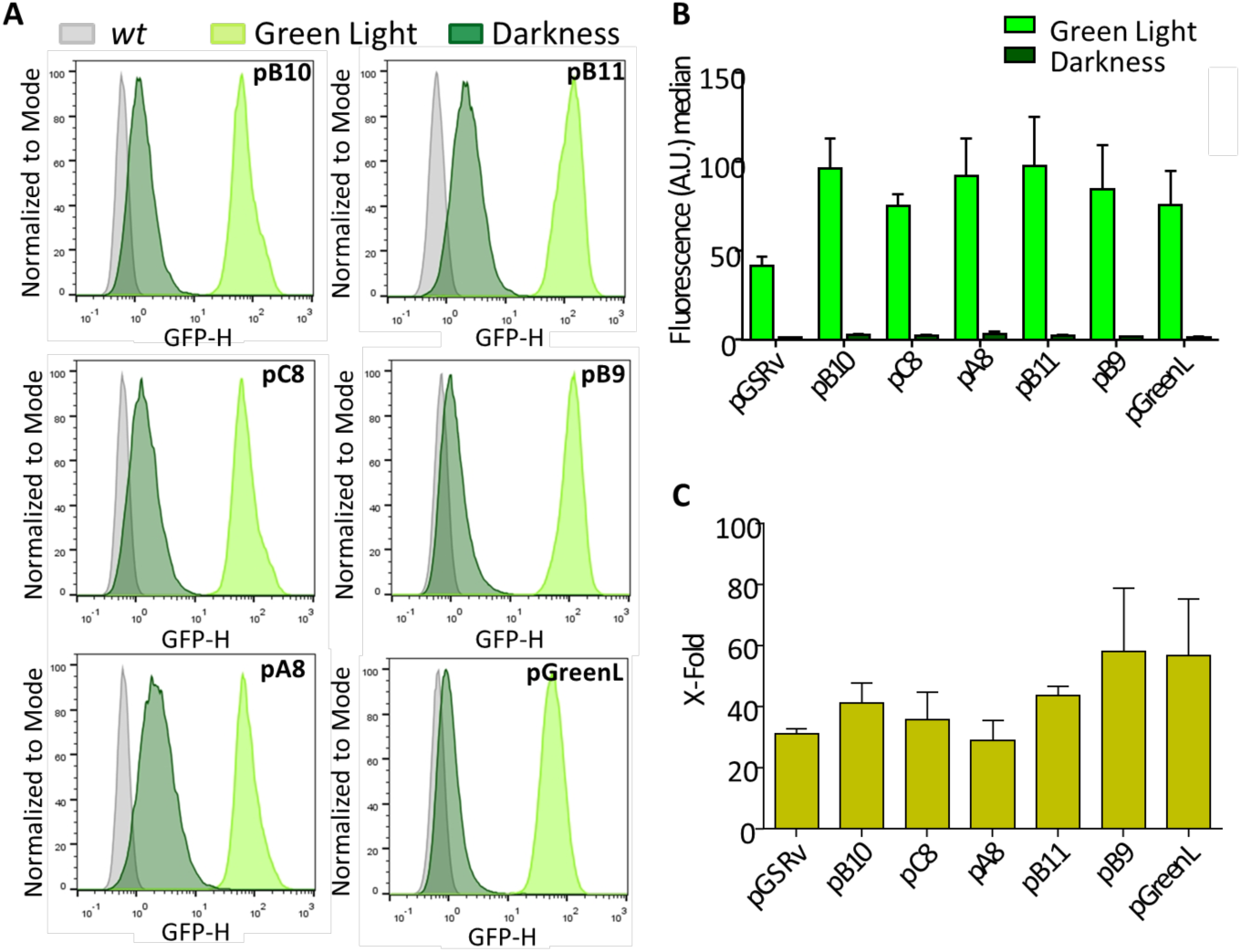
Light induction of the 5 best clones after the screening process. **A**, Cell cytometry for the 5 pGSRv derivatives carried by *P. putida*. Measurements include light-induced cultures (light green), darkness cultures (dark green) and the control *P. putida* KT2440 *wt* (grey). **B**. Median averages from three biological replicates of cell cytometry with induced and non-induced cultures, including the non-mutated plasmid pGSRv. **C**. Average X-Fold induction for each of the clones compared to pGSRv.

### Mapping mutations enabling inter-operativity of the CcaSR system

The next logic step was figuring out the evolutionary landscape of changes through the intergenic regions of the CcaSR system that had allowed a well-regulated performance in *P. putida*. To this end, the entire DNA constructs were subject to Pacific Biosciences SMRT (Single Molecule Real-Time) PCR amplicon sequencing, based on a modified version of Nyerges *et al*.^*12,23,24*^, as explained in Materials and Methods. The 5 analyzed samples included libraries Rv-A (sample 1) and Rv-B (sample 2), sorted libraries Rv-A (sample 3) and Rv-B (sample 4) and the 86 best performing clones (sample 5; Fig. 6). As expected, we found that mutations gathered just in the tagged regulatory regions. Non-sorted libraries RvA and RvB accumulated mutations in the sites that control expression of the two main proteins CcaS and CcaR along with others in the regulatory region of the *ho1-pcyA* genes. In contrast *P*_*cpcG2-172*_ included a lower number of changes. However, after the sorting process, we observed a different pattern of mutations in both libraries comparing them with the previous non-selected populations. The amount of mutations spotted in the regulatory region of CcaS increased notably —unlike the number of mutations included upstream of the rest of the genes. This finding suggested that regulation of CcaS expression was one of the key factors that influenced operation of the system. In the last sample (composed by the 86 best performing clones), we observed a similar distribution of mutations, confirming the relevance of CcaS expression levels for optimal reusability of the device. In a subsequent step, we sequenced the entire CcaSR system from pGreenL by classical Sanger sequencing in order to identify the changes that influenced the observed modifications of the basal level and the wider dynamic range of the sensor. We found 9 alterations located in only two regions: the RBS region of the CcaS protein had accumulated 7 mutations, while the RBS region of CcaR had 2 changes (the complete sequences can be found in the Supplementary files). The fact that the selected clone had only mutations in those regions strengthened our earlier hypothesis that the key bottleneck for re-nesting the optogenetic device from *E. coli* to *P. putida* was the imbalance of expression of main components of the CcaSR system. This evidence was also supported by the results of the RBS strength calculation given by Salis software^*25*^ (Supplementary Table S9), as the translation rates for *ccaS* and *ccaR* genes were re-balanced in the pGreenL clone.

**Figure 6:**
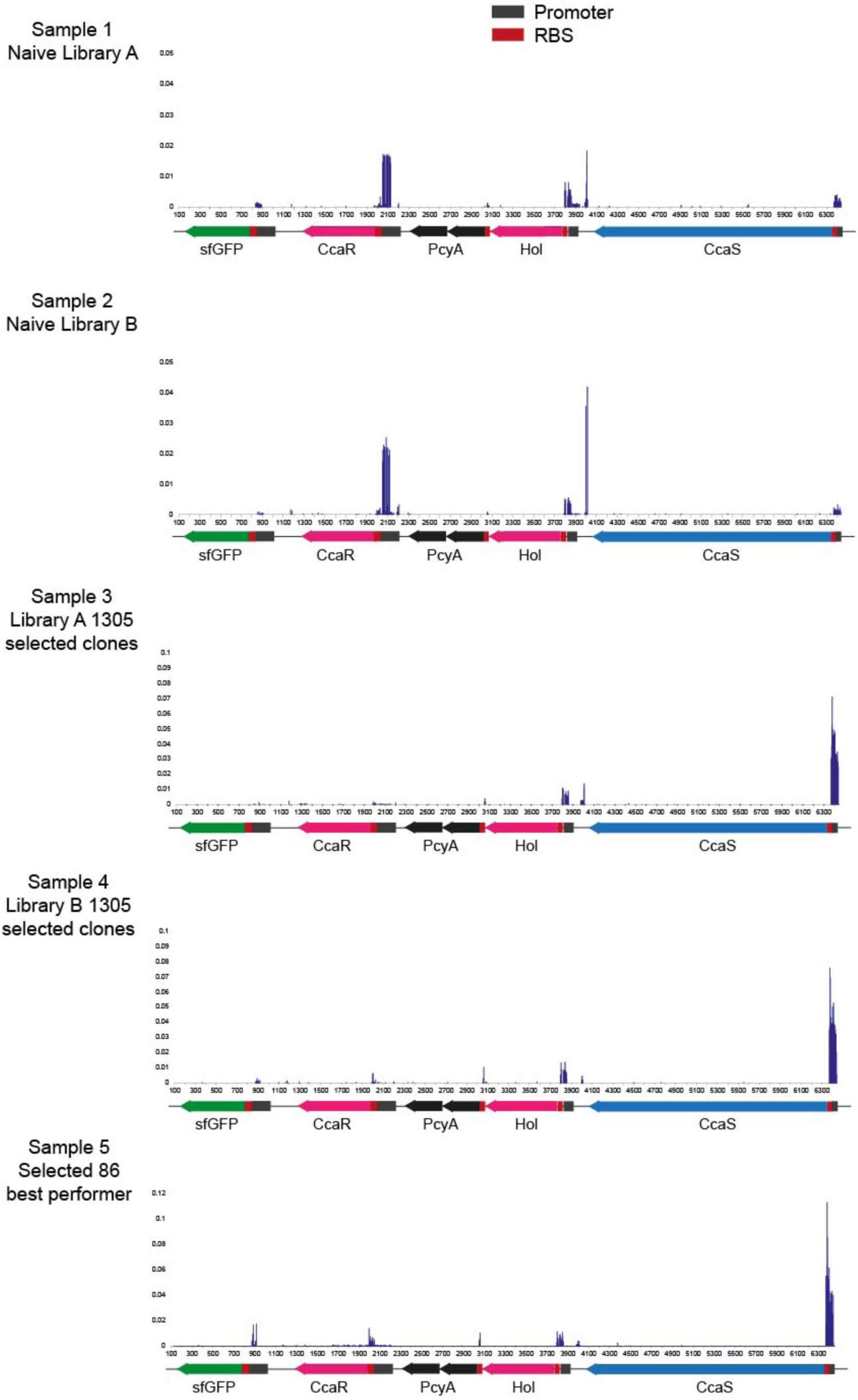
Deep sequencing of resulting libraries from the diversification process. Samples 1 and 2 correspond to naïve libraries RvA and RvB, both result of the mutagenesis process before sorting. Sample 3 and 4 correspond to the 1305 clones of library RvA and RvB (respectively) that were sorted and isolated for its characterization in terms of light responsiveness. Sample 5 represents the sequences of the 86 best performers that were selected from the 2610 (1305 + 1305) clones from samples 3 and 4.

### Conclusion

The functioning of any given genetic device depends not only on the intrinsic qualities of the biological parts involved or the relational logic among them. What ultimately makes them to work is the embodiment of a particular set of parameters that fit every transfer function to the preceding one and the next one for an optimal signal processing flow through the complete system^*26, 27*^. While design of electric circuits can be managed by adopting digital 0/1 abstractions, biological counterparts are complicated by the large number of parameters that affect the behavior of genetic constructs. Even when one engineered device is optimized for a desired performance in a certain host and fixed conditions, its conduct can typically change when moved to a different context—whether genomic, nutritional or physicochemical—let alone to other hosts^*28-30*^. The number of parameters to consider for such *interoperability* is so high that relocation of a genetic device from one biological recipient to another cannot be intended from first principles. Instead, the most useful approach is to set the system out to explore an ample solution space in the new environment and let it fluctuate while artificially rewarding displacement towards a preset optimum. The most popular approaches to this end involve combinatorial assembly of the biological parts involved^*9*^, with a focus on regulatory sequences (e.g. promoters and RBS sequences) and then enrichment or selection of the best combinations. While these stratagems have generally worked well, they ask for intensive labor (each diversification round requires extraction and manipulation of DNA) and the outcome is limited by the competence of the hosts to acquire exogenous DNA. As shown in the work above, most of these drawbacks can be overcome by adoption of DIvERGE^*12*^ a variant of MAGE (*Multiplex Automated Genome Engineering*^*31*^) technology that focuses the generation of hyper-diversity in specific sites of a target DNA segment *in vivo*. In the instance described in this work, we have exploited such a technique for simultaneously increasing mutation rates of the 5 regulatory regions of the light-responsive device [*ccaS•pcyA/ho1•cca*R•*P*_*cpcG2-172*_ →] of Fig. 3 with the objective of setting parameters suitable for its use in the soil bacterium *P. putida*. In our case, we have applied the technique to a device assembled in a low copy number plasmid in order to create libraries with a wide landscape of values and containing workable combinations for their use in the new host. As shown above, by first reducing the population with cell sorting and then isolating specific clones with a robot we ultimately found a new variant of the module with a highly improved performance. To the best of our knowledge, the resulting module [*ccaS•pcyA/ho1•cca*R•*P*_*cpcG2-172*_ →] adapted for use in *P. putida* constitutes the first optogenetic device available for this biotechnologically important bacterium^*10, 32, 33*^. Furthermore, the soft randomization-sorting-isolation-selection workflow hereby described should enable the portability and reusability of many other interesting devices initially developed for *E. coli* in other bacteria but the biotechnological value of which may be deployed only when passed to other species.

## Materials and Methods

### Culture conditions

For general cultures, *E. coli* and *P. putida* strains were grown in Luria Bertani (LB) medium, which had been filtered when performing Flow Cytometry experiments. Solid plates contained LB solidified with agar at 1.5% (w/v). Terrific Broth (TB) was used for the recovering of bacteria after oligo transformation during diversification. When necessary, antibiotics were added at the proper concentrations: kanamycin (Km, 50 µg mL^−1^), streptomycin (Sm, 50 µg mL^-1^) and gentamycin (Gm, 10 µg mL^-1^). For *E. coli*, the temperature used for incubation was 37°C, except with *E. coli* MG1655 strain bearing the mutagenesis machinery encoded in the pORTMAGE3 (Addgene Plasmid Number 72678, http://n2t.net/addgene:72678; RRID: Addgene_72678). In that case, incubation at 30°C was used for periods of non-induction and 42°C temperature was used for the induction of the mutagenesis apparatus. *P. putida* was incubated at 30 °C.

### Strains

Bacterial strains used in this study are described in Supplementary Table S4. For the construction of new plasmids, *E. coli* strains DH5α, JM109 and DH10B^+^ were used. For DIvERGE diversification, *E. coli* K12 MG1655 strain was used. *P. putida* KT2440^*34*^ was the strain selected for the final sorting and testing of the clones.

### Green light switch constructs

For most of the constructs, classical protocols of digestion, ligation and transformation in *E. coli* were used^*35*^. PCRs for new constructs were performed using Q5 High Fidelity polymerase (New England Biolabs, Ipswich, Massachusetts, USA), and analytic PCRs were performed using DNA AmpliTools Green Master Mix (BioTools, Madrid, Spain). Restriction enzymes were purchased from New England Biolabs (Ipswich, MA, USA) and T4 ligase (Roche, Basilea, Switzerland) was used for ligation purposes. Plasmid DNA was extracted from bacteria using QIAprep Spin Miniprep Kit (Qiagen, Venlo, Netherlands). Isolation of DNA from agarose bands was carried out using the NucleoSpin Extract II kit (Macherey Nagel, Düren, Germany). For the transformation of *E. coli* strains, either heat shock (90 seconds at 42 °C) or electrical shock (cuvettes 0.2 cm, 2.4 kv) was performed. For Gibson assembly reactions, both home-made mix^*36*^ and Gibson Assembly Master Mix were used (New England Biolabs, Ipswich, Massachusetts, USA). The complete list of constructed and used plasmids and primers can be found in Supplementary Tables S5 and S6. Plasmids pSR43.6 and pSR58.6 were kindly provided by J. Tabor and S. Schmidl^*5*^. They were used as templates for the amplification of the cassettes *CcaS-ho1-pcyA* and *CcaR-* P*cpc*G2-172-*msfGFP*, respectively. Gibson assembly reaction was performed for one side using the *CcaS-ho1-pcyA* cassette and pSEVA631 (digested with EcoRI and HindIII). Another Gibson assembly reaction was performed to connect the *CcaR-*P_*cpc*G2-172_ -*msfGFP* cassette and pSEVA241 (digested with EcoRI and HindIII again). The resulting respective plasmids, pCcaSPCB and pCcaRGFP, were altogether transformed into *E. coli* JM109 by electroporation. Accuracy of the cloned DNA fragment was confirmed by DNA sequencing. After that, they were transformed into *P. putida* KT2440, wherein their function was tested. In a second batch of constructs, pGSFw and pGSRv plasmids were built assembling the *CcaS-ho1-pcyA* and *CcaR-*P_*cpc*G2-172_ -*msfGFP* cassettes, previously amplified using oligos listed in the Supplementary files, with a PacI/HindIII digested pSEVA621 backbone. Gibson reaction was performed for the assembly. pGSFw and pGSRv were transformed into *E. coli* MG1655 for diversification. The outcome libraries and the parental non mutated plasmids were later transformed into *P. putida* as described previously. After diversification, characterization and sequencing of the selected clone, the optimized light switch cassette was re-cloned in a pSEVA621 using AscI and SwaI enzymes. The final plasmid was called pGreenL and it was characterized as explained above.

### DIY Arduino-controlled light panel

For illumination with green light, we used a computer that controlled an RGB LED panel matrix connected to an Arduino device. Components used for the assembly of the LED panel and Arduino are listed in Supplementary Table S1. For the Arduino UNO board, a firmware was developed in the Arduino IDE v1.0.5r2. The code also needed two libraries provided by the manufacturer of the LED panel: <Adafruit_GFX.h> found at https://github.com/adafruit/Adafruit-GFX-Library and <RGBmatrixPanel.h> found at https://github.com/adafruit/RGB-matrix-Panel. Arduino board firmware waits for user commands from the PC through a USB-CDC interface. When commands are received, it checks if they have a correct format and then proceeds to the colour and shape update in the LED matrix panel. Although the Arduino-LedMatrix composite can show different shapes and colours, green and red were the only configuration used in all the LEDs of the panel. Once the LED panel matrix is setup, the user can send a command to save the current colour and shape. Therefore, the next time the Arduino device is powered up there is no need to reprogram it again. The firmware code was developed using a subset of C/C++ language with the Arduino Native libraries. The Arduino UNO device was connected to the LED panel. Power supply connectors were also joined between Arduino and the panel. A specific software was designed to let the user write commands about the light colour and the number of switched-on LEDs. Then, commands are sent to the Arduino board through the USB connector. Once the Arduino board processes them, the colours and shapes in the LED matrix panel are updated. This code is a Python 2.7.8 script for Windows 7, and it also uses the python libraries pyserial-2.7, pywinusb-0.3.3 and, setuptools-5.4.2. Once the Arduino board is programmed, the connection to the PC can be removed so that both the Arduino board and the LED matrix panel composite are portable. Disposition of LED panel, Arduino board and computer for the illumination of cultures is shown in Supplementary Fig. S2.

### Diversification

Directed Evolution with Random Genomic Mutations (DIvERGE) experiments were performed using the procedure described by Nyerges and his colleagues^*12*^. Oligos, designed for DIvERGE, were synthesised at the Nucleic Acid Synthesis Laboratory of the Biological Research Centre, Hungarian Academy of Sciences (Szeged, Hungary). Soft-randomized oligos (listed in the Supplementary Table S7) typically included 1 to 25 mutations in the desired region that can later be introduced to the targeted region of the genome (or a plasmid) of the microorganism of interest. *E. coli* MG1655 strain was cotransformed with pORTMAGE3 and pGSFw plasmids for one side, and the same strain was cotransformed with pORTMAGE3 and pGSRv for the other side. As a result, two new MG1655 strains (named Fw and Rv) were constructed carrying the machinery for DIvERGE and the cassette *CcaS-ho1-pcyA-CcaR-*P_*cpc*G2-172_ -*msfGFP* in the forward orientation (Fw strain) and in the reverse orientation (Rv strain). Both cultures, carrying either the pGSFw or the pGSRv plasmid, were in turn exposed to 5 rounds of DIvERGE mutagenesis using two combinations of oligos per each culture: 1+2+3+4+5a which produced a library A, and 1+2+3+4+5b which produced a library B (Supplementary Table S2). The difference between oligo 5a and 5b is that the former does not target the G box of the p_*cpc*G2-172_ regulatory region where CcaR binds^*4,5*^ and, in consequence, does not introduce mutations, while oligo 5b does target this region. As a result, we had four different libraries coming from the four steps of diversification of two plasmids.

### Transformation of *P. putida* and cell sorting enrichment

*E. coli* MG1655 bearing the libraries Fw-A, Fw-B, Rv-A and Rv-B were cultured separately overnight (O/N) in 20 mL of LB medium. Afterwards, DNA plasmid from the total 20 mL was extracted and transferred into *P. putida* via 4 different transformations per library. Cultures were incubated O/N in liquid media with the appropriate antibiotic, and then subject to the first screening. First, 100 µL of the saturated cultures were taken to inoculate 10 mL of LB in a flask, and they were incubated under green light conditions during 16 h. The rest of the *P. putida* library was preserved in a glycerol stock. After the light incubation, 1 mL of the culture was taken, and the OD_600_ was adjusted to 0.5. Then, the different libraries were sorted using a cell sorter FACSVantage SE with an installed BD FACSDiVa Software (Becton Dickinson, Franklin Lakes, New Jersey, USA). The top GFP fluorescent population was selected in every sorting event (Supplementary Fig. S3), while the rest of the clones were discarded. Final volume of the outcome was supplemented to reach a total of 10 mL and was then grown O/N. Next, 100 µL were used to inoculate another 10 mL LB media for a new cycle of green light exposition and cell sorting, and the rest of the culture was saved in glycerol stock at −80°C. After three sorting cycles that included incubation under green light during 16 h followed by selection of the most promising GFP positive clones and the preservation of the outcome, the population containing the most green fluorescent cells was incubated in complete darkness for 16 h and passed through a final sorting with the opposite selection criteria, that being selection of the cells with a lower fluorescence. The final outcome was incubated O/N at 30 °C and preserved in glycerol at −80 °C.

### Isolation of sorted clones

For the following steps, only the RvA and RvB libraries were used, as they showed a higher variability in terms of response to light. For the colony picker screening, we used 30×30 cm plates with 300 mL LB agar. O/N cultures of the RvA and RvB libraries obtained from the sorting cycles described above were diluted 1 to 50 in LB, and 100 µL of the outcome were spread in each square plate, to plate the maximum amount of colonies with an adequate separation amongst them. Plates grown O/N at 30°C were placed in a colony picker Getenetix QPix (Molecular Devices, San José, California) that took and inoculated clones in 96-well microtiter plates containing 200 µL of LB in each well. In total, 15 microtiter plates harboring 1305 variants from the RvA library and another 15 with the same amount of clones from the RvB library were obtained by this procedure and preserved by addition of 15% glycerol (Supplementary Fig. S4). Each of those 30 plates keeping the 2610 selected clones was organized as indicated in Supplementary Fig. S4A. Supplementary Fig. S4B shows the workflow from the sorted libraries, to the colony picker isolation and finally to the selected clone.

### Selection of best performers

Each plate containing the isolated clones was used to inoculate 200-µl aliquots placed on two replica 96-well plates for fluorescence measurement (Costar black plates with clear bottom; Thermo Fisher Scientific Inc., Pittsburgh, PA, USA; Supplementary Fig. S4). One of the replica plates was placed under darkness conditions (covered with aluminium foil) while the other under light conditions during 16 h without shaking. Both plates were measured for OD_600_ and fluorescence using a SpectraMax Plate reader (Molecular Devices, San José, CA, USA). After normalisation of GFP values using the OD_600_, X-fold induction was calculated for each clone. Mean, standard error (SD) and covariance (CV) were calculated for the X-Fold of the 8 replicates of the parental strain. CV illustrates the reproducibility of the parental strain results; this variability probably extends as well to the results of each clone. Each calculated X-Fold for the isolated clones was divided by the X-Fold mean of the parental strain. If the resulting number, deemed the relative X-Fold, was superior to 1+ CV/100, it was considered to be a significant value. The higher this value was, the better the clone in regards to its induction capacity. After analyzing all the isolated variants, the best 86 performers were further examined in three biological replicates using the same method. Finally, the five clones that showed the best behavior were studied by single cell flow cytometry experiments in more detail.

### Flow cytometry

Overnight cultures of each of the five best performers were used to inoculate fresh filtered LB (1:5000) in 10 mL tubes and incubated with 190 rpm shaking at 30°C during 8 hours in duplicate. One of the replica was grown in darkness (covered with aluminium foil) and the other in a light-induced state. Next, cells were sedimented and washed with filtered 1X PBS and resuspended in the same buffer at the adequate OD_600_ to pass them through a MACSQuant Flow Citometer (Miltenyi, Bergisch Galdbach, Germany). An Ar laser, diode-pumped solid state, was used to excite msfGFP at 488 nm and the fluorescence signal was recovered with a 525/40 nm band-pass filter. For the experiments, ≥120,000 events were counted per sample. Results were analysed using FlowJo software (FlowJo LLC, Ashland, Oregon).

### Deep amplicon sequencing of DIvERGE libraries and individual clones

The allelic composition of the CcaSR libraries and the best-performer ten clones were determined by a Pacific Biosciences SMRT (Single Molecule Real-Time) PCR amplicon sequencing protocol^*12*^. To create sequencing libraries, a 6595 bp long region, encoding *ccaS, pcyA, ho1, ccaR*, P_*cpc*G2-172_ -*msfgfp*, was PCR amplified from the previously isolated, pooled plasmid DNA samples by using the corresponding barcoded primer pairs (see Supplementary Table). To multiplex sequencing samples, symmetrically barcoded PCR primers were designed^*12*^. Each primer-pair consisted of a symmetrical 16 nucleotide-long Pacific Biosciences Sequel barcode sequence, a 3 nucleotide-long 5’ terminal sequence, besides the 3’ terminal plasmid-specific primer sequences. All PCR primers were synthesized by the Nucleic Acid Synthesis Laboratory of the Biological Research Centre, Hungarian Academy of Sciences, Szeged, Hungary, and purified using high-performance liquid chromatography (HPLC). Following DNA synthesis and purification, primers were suspended in nuclease-free H2O at 100 µM final concentration. Next, the [*ccaS•pcyA/ho1•cca*R• *P*_*cpcG2-172*_→] region from each plasmid sample was amplified in 3×25 µl volumes, consisting of 37.5 µl 2× Q5 Hot-Start MasterMix (New England Biolabs), 2.5 µl of the corresponding sample-specific, barcoded primers (10 µM), 200 ng template plasmid DNA, and 33 µl nuclease-free H2O. PCRs were performed in thin-wall PCR tubes in a BioRad CFX96 qPCR machine with the following thermal profile: 98 °C 3 minutes, 20 cycles of (98 °C 15 seconds, 72 °C 4:30 minutes), and a final extension of 72 °C for 10 minutes. Following PCRs, the 6633 basepair-long amplicons were purified by using a Zymo Research DNA Clean and Concentrator™ Kit according to the manufacturer’s protocol (Zymo Research) and eluted in 30 µl 0.5× Tris-EDTA (TE) buffer (pH 8.0). To prepare samples for sequencing, amplicons were quantified using Qubit dsDNA BR assay kit (Thermo Fisher Scientific), mixed, and libraries were sequenced on a Pacific Biosciences Sequel instrument using Sequel Polymerase v2.1 on an SMRT cell v2 with Sequencing chemistry v2.1 (both from Pacific Biosciences). The SMRT cell was loaded by diffusion, and the sequencing movie time was 1200 min. After sequencing, raw sequencing subreads were demultiplexed according to their corresponding barcodes (see Supplementary Table S8) by using Demultiplex Barcodes pipeline on SMRT Link v5.1.0.26412 (SMRT Tools v5.1.0.26366). A minimum barcode score of 26 was used identify high-quality barcodes. The average sequencing read counts were 9000 per library sample. To determine the sequence of individual clones, we relied on Long Amplicon Analysis and determined the phased consensus sequences for each of the corresponding demultiplexed datasets. Following demultiplexing, sequencing reads were mapped to their corresponding reference sequence by using bowtie2 2.3.4^*37*^ in “--very-sensitive” mode and the nucleotide composition was extracted for each nucleotide position within the CcaSR region. Finally, allelic replacement frequencies at nucleotide positions were quantified by measuring the distribution and ratio of nucleotide substitutions for each reference nucleotide position.

## ASSOCIATED CONTENT

**Supplementary Figure S1**. Representation of the CcaSR two-component system and its mechanism of action.

**Suplementary Figure S2**. LED matrix panel used to induce the CcaSR system with green light.

**Supplementary Figure S3**. Schematic representation of the procedure followed for the libraries’ cell sorting.

**Figure S4**. Isolation of clones from libraries.

**Supplementary Figure S5**. Outcome plates at different points of the screening.

**Supplementary Figure S6:** pGreenL behaviour. **Supplementary Table S1**. List of components used for the LED plate and the company where they were purchased.

**Supplementary Table S2**. Plasmids and oligos used for DIVERGE diversification and the resulting libraries.

**Supplementary Table S3**. Sorting rounds of each library and their selection events.

**Supplementary Table S4**. List of strains used in this study

**Supplementary Table S5**. List of plasmids used in this study

**Supplementary Table S6**. List of primers used for the constructions performed in this study.

**Supplementary Table S7**. List of primers used for the diversification of the regulatory regions.

**Supplementary Table S8**. List of primers used for the sequencing for the mutations profile.

**Supplementary Table S9**. Results of the RBS strength calculation for the original (WT) and the mutated (pGreenL) RBS regions of *ccaS* and *ccaR* genes using Salis software.

**pGreenL sequence**

## AUTHORS’ CONTRIBUTIONS

AN and VDL conceived the study and BC, AHG and AN designed the experiments. AHG carried out the experimental parts of the work. AHG, VDL and AN wrote the article. All the authors contributed to the discussion of the research and interpretation of the data.

## ACKNOWLEDGEMENTS

Authors are indebted to Laura Molero from the SIdI service at the Universidad Autónoma de Madrid for the sorting of libraries. We also thank David Gonzalez and Miguel Alcalde for their help with the isolation of colonies using their Colony Picker QPIX platform. We also express our gratitude to Raúl Rodríguez-Barreiro for his assistance with the led plate assembly. This work was funded SETH Project of the Spanish Ministry of Science RTI 2018-095584-B-C42, MADONNA (H2020-FET-OPEN-RIA-2017-1-766975), BioRoboost (H2020-NMBP-BIO-CSA-2018), and SYNBIO4FLAV (H2020-NMBP/0500) Contracts of the European Union and the S2017/BMD-3691 InGEMICS-CM of the Comunidad de Madrid (European Structural and Investment Funds). Sequencing services were provided by the Norwegian Sequencing Centre (Research Council of Norway and the Southeastern Regional Health Authorities). CP was supported by contracts H2020-ERC-2014-CoG 648364, the Wellcome Trust, GINOP MolMedEx TUMORDNS, GINOP-2.3.2-15-2016-00020, GINOP (EVOMER) GINOP-2.3.2-15-2016-00014 and the Lendület Program of the Hungarian Academy of Sciences. AN was the recipient of a a PhD fellowship from the Boehringer Ingelheim Fonds.

