## Supplementary for "Multiple-site diversification of regulatory sequences enables inter-species operability of genetic devices"

### SUPPLEMENTARY MATERIALS

**Figure S1.** Representation of the CcaSR two-component system and its mechanism of action.

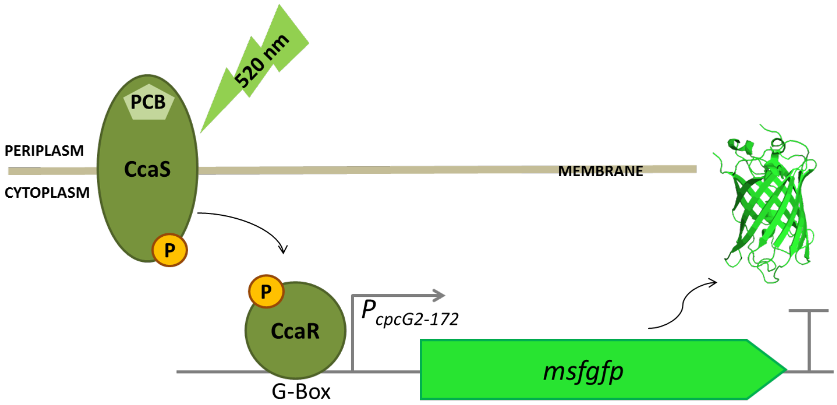


In the presence of cofactor phycocyanobilin (PCB), CcaS senses green light (520 nm) and gets autophosphorylated. In that form, it is able to pass the phosphoryl group to the transcription factor CcaR. In the phosphorylated state, CcaR recognizes the G-Box of the promoter $P_{cpcG2\text{-}172}$, recruits the $\sigma^{70}$ containing RNA polymerase and activates the transcription of the downstream gene, which in this case is the monomeric superfolder version of GFP[7, 9].

**Figure S2:** LED matrix panel used to induce the CcaSR system with green light.

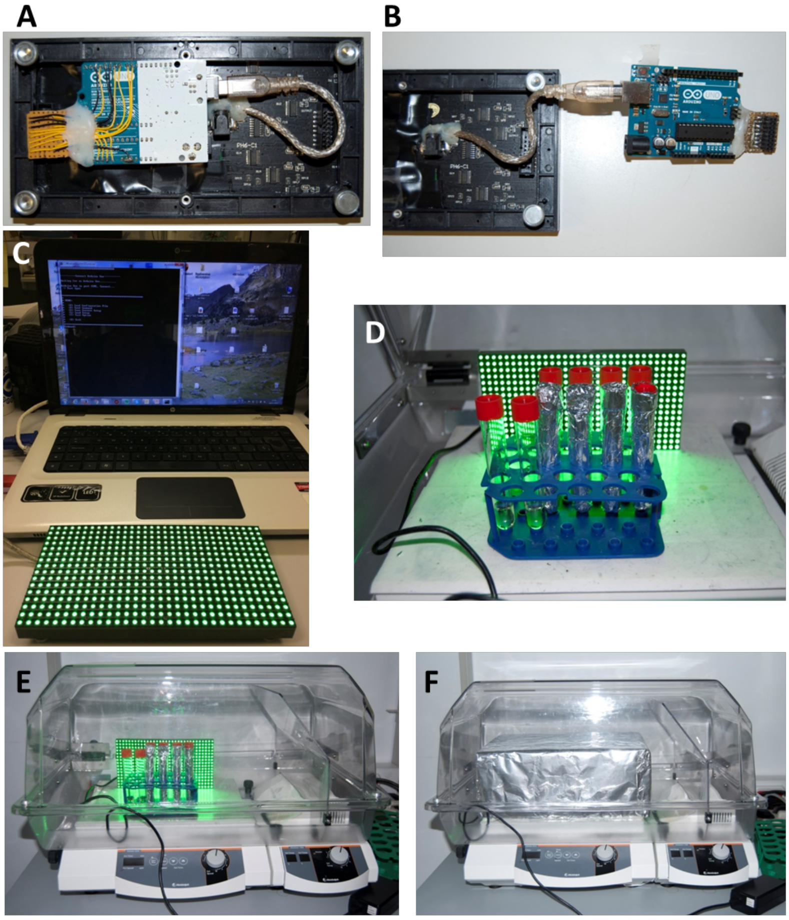


**A** and **B.** LED matrix panel and Arduino UNO device connexions with the power supply cable. **C.** Computer software controlling panel illumination. **D.** Tubes under green light induction and in darkness (covered with aluminium foil) inside shaker incubator. **E.** General view of the incubator and illumination conditions. **F.** Box covering cultures to avoid an excess of white environmental light entering into tubes that were exposed to green light.

**Figure S3:** Schematic representation of the procedure followed for the libraries' cell sorting**.**

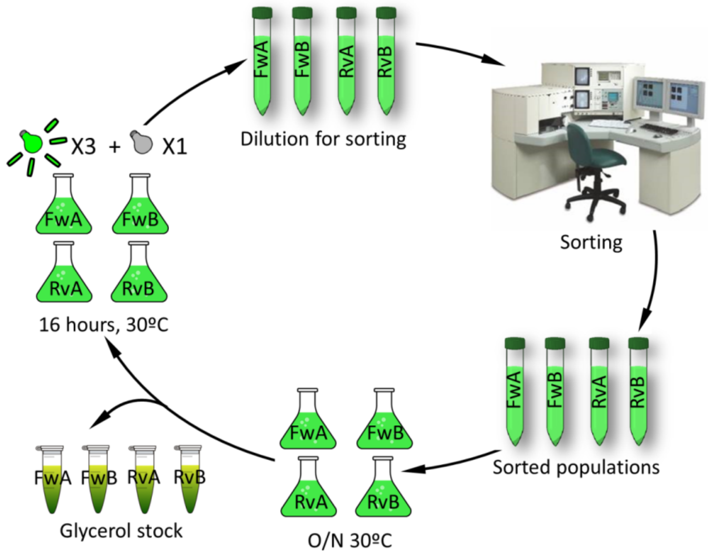


**Figure S4:** Isolation of clones from libraries.

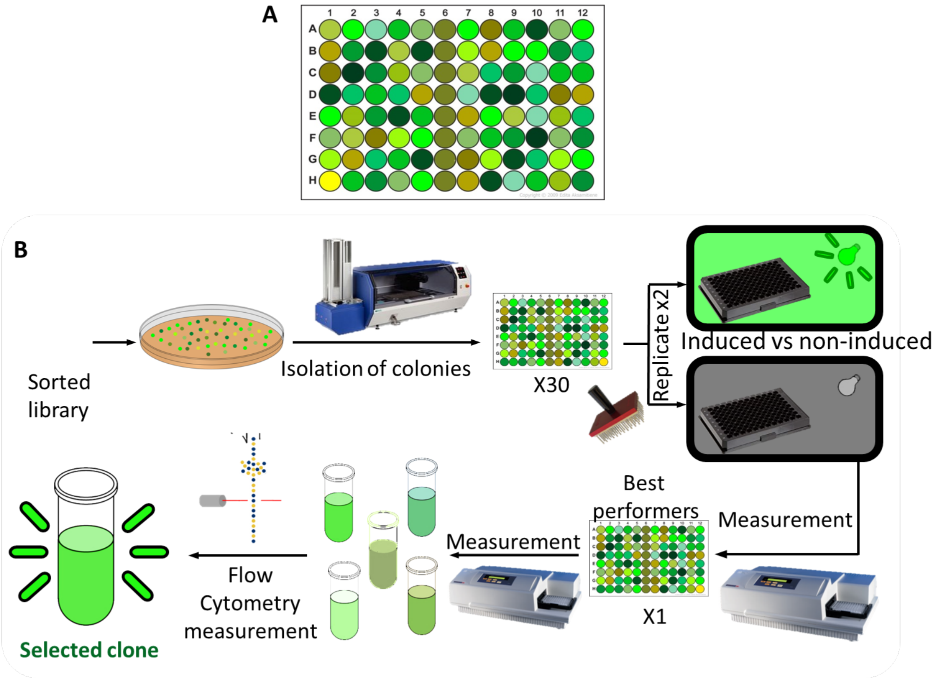


**A.** An example distribution for the 96 microtiter plates that were cultured under light induction or under darkness for the measurement of each clone. Column 6 was only inoculated with *P. putida* bearing the wild type parental plasmid pGSRv. Well H1 was used as a blank into which 200 µL of LB was added without inoculation. **B.** Sketch of the screening procedure described in sections 4.5 and 4.6 starting with *P. putida* harbouring a sorted library and ending with a final clone being selected as the best performer.

**Figure S5: Outcome plates at different points of the screening.**

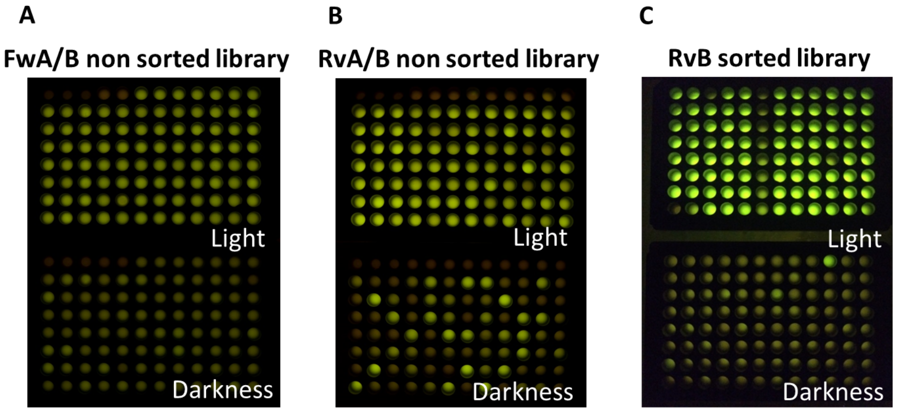


Isolated clones from non-sorted Rv **A.** and Fw **B.** libraries showed different diversity rates in response to light. In both Rv and Fw libraries, the 96-well plate at the top was induced by light and the plate at the bottom was an exact replicate kept in darkness. A higher variability in GFP expression can be observed in Rv libraries, suggesting that the mutagenesis was more successful in pGSRv plasmid. Notice that in these two pairs of plates, column 6 was not inoculated with the parental plasmid pGSRv as other mutated variants were instead grown in those wells. **C.** Analysis of Rv B/ pGSRv-derived clones. The sorted library exhibited in general a better OFF/ON ratio. Note the difference in the behaviour of wells in column 6 (this time inoculated with *P. putida* bearing non-mutated pGSRv) and that of the rest of the clones, as our process selected the best-behaving population.

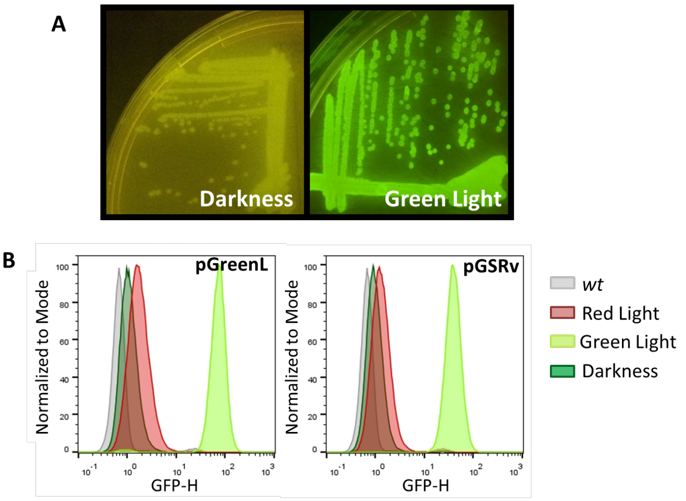


**Figure S6:** pGreenL behaviour.

**A.** Picture of LB agar streak of *P. putida* carrying pGreenL induced by green light or kept in darkness. **B.** Low GFP production when cultures were illuminated with red light (red) proved that the light sensor was specifically induced with green light (light green). In the graph, darkness non-induced cultures (dark green) and the control (grey) are also plotted.

**Table S1.** List of components used for the LED plate and the company where they were purchased.

| **Product** | **Company** | **Cat. number** |
|---|---|---|
| Medium 16x32 RGB LED matrix panel | Adafruit Industries (New York City, New York, USA) | 420 |
| Board, Arduino Uno, MCU ATmega328P, 14 E/S 3.3V, 6 analogic inputs, 6 outputs PWM | Farnell (Leeds, United Kingdom) | A000066 |
| Samtec SSM-108-L-DV receptacle 2,45 mm, SMT, 16 routes | Farnell (Leeds, United Kingdom) | 2308487 |
| Ideal Power 25 HK-AB-050A400-D5-1 Power Supply, Ext PLG-IN, $4^{a}$, 5V, 20W | Farnell (Leeds, United Kingdom) | 2334612 |
| PRO ELEC PL00374 power cable, Euro to Iec, 2M, 10A | Farnell (Leeds, United Kingdom) | 3507737 |
| Lumberg 161314 chasis socket, PSU, SMD, 3.5A | Farnell (Leeds, United Kingdom) | 1243245 |
| Switchcraft PC722A Jack Socket, DC | Farnell (Leeds, United Kingdom) | 1608744 |

**Table S2.** Plasmids and oligos used for DIVERGE diversification and the resulting libraries.

| **Original diversified plasmid** | **Pool DIvERGE oligos** | **Resulting library** |
|---|---|---|
| pGSFw | A: 1 + 2 + 3 + 4 + 5a | Fw A |
| pGSFw | B: 1 + 2 + 3 + 4 + 5b | Fw B |
| pGSRv | A: 1 + 2 + 3 + 4 + 5a | Rv A |
| pGSRv | B: 1 + 2 + 3 + 4 + 5b | Rv B |

**Table S3.** Sorting rounds of each library and their selection events.

| | **Library Fw A** | **Library Fw B** | **Library Rv A** | **Library Rv B** |
|---|---|---|---|---|
| **Selected events 1$^{st}$ round** | 20 552 | 21 651 | 28 840 | 22 072 |
| **% 1$^{st}$ round** | 11.5 | 9.9 | 5.6 | 5.2 |
| **Selected events 2$^{nd}$ round** | 25 406 | 27 565 | 26 119 | 25 334 |
| **% 2$^{nd}$ round** | 9.9 | 9.0 | 1.7 | 1.4 |
| **Selected events 3$^{rd}$ round** | 86 567 | 82 220 | 215 155 | 85 662 |
| **% 3$^{rd}$ round** | 46.4 | 36.3 | 44.7 | 59.8 |
| **Selected events negative round** | 50 268 | 30 238 | 47 450 | 51 152 |
| **% negative round** | 2.2 | 2.2 | 48.3 | 45.6 |

**Table S4. List of strains used in this study**

| Strain | Description | Reference |
|---|---|---|
| DH5α | $F^-$, *supE44*, Δ*lacU169*, (φ80 lacZDM15), *hsdR17*, (*rkmk*$^+$), *recA1*, *endA1*, *thi1*, *gyrA*, *relA* | 1 |
| JM109 | F′, *traD36*, *proA+B+*, *lacIq*, "(*lacZ*)M15/ "(*lac-proAB*), *glnV44* e14-, *gyrA96*, *recA1*, *relA1*, *endA1*, *thi*, *hsdR17* | 2 |
| DH10B+ | $F^-$, mcrA, Δ(*mrr-hsd*RMS-*mcr*BC), Φ80d*lac*ZΔM15, Δ*lac*X74, *end*A1, *rec*A1, *deo*R, Δ(*ara*,*leu*)7697, *ara*D139, *gal*U, *gal*K, *nup*G, (Str$^R$), *rps*L $\lambda^-$ | 3 |
| MG1655 | K12 MG1655 wild type, F-, *lambda-*, *rph-1* | 4 |
| KT2440 | Prototrophic, wild-type strain derived of *P. putida* mt-2 without pWWO plasmid | 5 |

**Table S5: List of plasmids used in this study**

| Plasmid | Description | Reference |
|---|---|---|
| pSEVA631 | GmR, oriV, oripBBR1, oriT, standard broad host range | 6 |
| pSEVA241 | KmR, oriV, oriColE1, oriT, standard broad host range | 6 |
| pSEVA621 | GmR, oriV, oriRK2, oriT, standard broad host range | 6 |
| pSR43.6 | SpecR, orip15A, *ccaS*, *ho1-pcyA* | 7 |
| pSR58.6 | CmR, oriColE1, *ccaR*, $p_{cpcG2\text{-}172}$ →*msfGFP* | 7 |
| pORTMAGE3 | λ-Red + strong-RBS-mutLE32K expressing MAGE vector, derived from pSIM8 (pBBR1 broad host-range origin of replication) | 8 |
| pCcaSPCB | pSEVA631 derivative, GmR, oripBBR1, *ccaS*, *ho-pcyA* | This work |
| pCcaRGFP | pSEVA241, KmR, oriColE1, *ccaR*, $p_{cpcG2\text{-}172}$-→*msfGFP* | This work |
| pGSFw | GmR, oriRK2, *ccaS*, *ho1-pcyA*, *ccaR*, $p_{cpcG2\text{-}172}$ →*msfGFP*, forward orientation | This work |
| pGSRv | GmR, oriRK2, *ccaS*, *ho1-pcyA*, *ccaR*, $p_{cpcG2\text{-}172}$ →*msfGFP*, reverse orientation | This work |
| pGreenL | pGSRv derivative, point mutations at regulatory regions for *ccaS* and *ccaR*, numbered previously as B9 clone | This work |

**Table S6:** List of primers used for the constructions performed in this study.

| Primer name | Sequence(5'→3') | TM | Plasmid |
|---|---|---|---|
| 43.6 F | AACAATTTCACACAGGAGGCCGCCTAGGCCGCGGCCGC GCCAAAAACCCCAGTTTTTACGG | 68 ºC | pCcaSP CB |
| 43.6 R | AGACTAGTCGCCAGGGTTTTCCCAGTCACGACGCGGCCG CTTATTGGATAACATCAAATAAGAC | 68 ºC | pCcaSP CB |
| 58.6 F | AACAATTTCACACAGGAGGCCGCCTAGGCCGCGGCCGC GCTTGACGGCTAGCTCAGTCCTA | 68 ºC | pCcaR GFP |
| 58.6 R | AGACTAGTCGCCAGGGTTTTCCCAGTCACGACGCGGCCG CTCATCATTTGTACAGTTCATCC | 68 ºC | pCcaR GFP |
| 43.6Fw uni F | GCCTTTCGTTTTATTTGATGCCTTTAATCCCAGTTTTTAC GGCTAGCTCAGTC | 70 ºC | pGSFw |
| 43.6Fw uni R | CTGTACCTAGGACTGAGCTAGCCGTCAACCTCATGCGAA ACGATCCTCATC | 70 ºC | pGSFw |
| 58.6Fw /Rv uni F | TTGACGGCTAGCTCAGTCCTAGGTACAG | 70 ºC | pGSFw / pGSRv |
| 58.6Fw uni R | GTTTTCCCAGTCACGACGCGGCCGCAAGCTTTCATCATT TGTACAGTTCATCCATACC | 70 ºC | pGSFw |
| 43.6 Rv uni F | GTTTTCCCAGTCACGACGCGGCCGCAAGCTTTCAAAAAC CCCAGTTTTTACGGC | 70 ºC | pGSRv |
| 43.6 Rv uni R | CTGTACCTAGGACTGAGCTAGCCGTCAATTACTTTGCAG GGCTTCCCAACC | 70 ºC | pGSRv |
| 58.6 Rv uni R | GCCTTTCGTTTTATTTGATGCCTTTAATTCATCATTTGTA CAGTTCATCCATACC | 70 ºC | pGSRv |
| PacBio F | CTAGGGCGGCGGATTTGTCC | 66 ºC | PacBio seq amplification |
| PacBio R | GCGGCAACCGAGCGTTCTG | 64 ºC | PacBio seq amplification |

**Table S7.** List of primers used for the diversification of the regulatory regions.

| Oligo ID | Sequence 5'-3' |
|---|---|
| | The underlined, italicized, bold regions are synthesized with 5% monomer soft-randomization |
| DIvERGE 1 | 5’-AAAACCCCAGTT***TTTACGGCTAGCTCAGTCCTAGGTATAGTGCTAGCTATAGAGGTTAAAATCTTAATAGGAGGAAGGCGCA***ATGGGCAA |
| DIvERGE 2 | 5’-ACCGTCAAAAAAA***CTGACAGCTAGCTCAGTCCTAGGTATAATGCTAGC***TCACACAGAATTCATT***AAAGAGGAGAAA***GGTACCATGAGTGT |
| DIvERGE 3 | 5’-AGTTGGCCTCGCCACCTCCGAAGGCTAGTT***AAAGAGGAGAAA***GGATCCATGGCCGTCACTGATTTAAGTTTGACCAATTCTTCCCTGAT |
| DIvERGE 4 | 5’-GCCGCGGCCGCGC***TTGACGGCTAGCTCAGTCCTAGGTACAGTGCTAGCTGTTAAACGCTCCTCGTAAGAACACGGGAAAGCA***ATGAGAAT |
| DIvERGE 5a | 5’-TTCTTTACGATTTT***CTCCCCCTTTTCTTCAATTTTACTTTGTTAGGATCGCATTTTTAAAAAGAGGAGAAATACTAG***ATGCGTAAAGGCG<br><br>Underlined+italicized regions are synthesized with 5-5-5% soft-randomized monomer mixture<br>***A*** **=** 85% A+5%C+5%T +5%G<br>***C*** **=** 85% C+5%A+5%T +5%G<br>***G*** = 85% G+5%C+5%T +5%A<br>***T*** = 85% T+5%C+5%A +5%G<br>The underlined, italicized, bold regions are synthesized with 0,5% monomer soft-randomization |
| DIvERGE 5b | 5’-***CAGTTTTAGTCGCATCAGCTAACTTTCCGATTTCTTTACGATTTTCTCCCCCTTTTCTTCAATTTTACTTTGTTAGGATCGCATTTTTAA***<br><br>Underlined+italicized regions are synthesized with 0,5 - 0,5 - 0,5% soft-randomized monomer mixture<br>***A*** **=** 98,5% A+0,5%C+0,5%T +0,5%G<br>***C*** **=** 98,5% C+0,5%A+0,5%T +0,5%G<br>***G*** = 98,5% G+0,5%C+0,5%T +0,5%A<br>***T*** = 98,5% T+0,5%C+0,5%A +0,5%G |

**Table S8.** List of primers used for the sequencing for the mutations profile.

| Sample ID | Primer ID | Primer sequence (5'-3') | Barcode sequence |
|---|---|---|---|
| **SEVA_LIB1 / Sample1** | **PABF1 051** | ATACGTGTCTAGCGCGCGCCTA GGGCGGCGGATTTGTCC | **CGTGTCTAG CGCGCGC** |
| | **PABR1 051** | AATGCGCGCGCTAGACACGGCG GCAACCGAGCGTTCTG | |
| **SEVA_LIB2 / Sample2** | **PABF1 052** | ATAGTGTGAGATATATATCCTAG GGCGGCGGATTTGTCC | **GTGTGAGAT ATATATC** |
| | **PABR1 052** | AATGATATATATCTCACACGCGG CAACCGAGCGTTCTG | |
| **SEVA_LIB3 / Sample3** | **PABF1 053** | ATACTCACGTACGTCACACCTAG GGCGGCGGATTTGTCC | **CTCACGTAC GTCACAC** |
| | **PABR1 053** | AATGTGTGACGTACGTGAGGCG GCAACCGAGCGTTCTG | |
| **SEVA_LIB4 / Sample4** | **PABF1 054** | ATAGCGCACGCACTACAGACTAG GGCGGCGGATTTGTCC | **GCGCACGCA CTACAGA** |
| | **PABR1 054** | AATTCTGTAGTGCGTGCGCGCG GCAACCGAGCGTTCTG | |
| **SEVA_LIB5 / Sample5** | **PABF1 055** | ATACACACGAGATCTCATCCTAG GGCGGCGGATTTGTCC | **CACACGAGA TCTCATC** |
| | **PABR1 055** | AATGATGAGATCTCGTGTGGCGG CAACCGAGCGTTCTG | |
| **SEVA_COLONY 1 / Sample6** | **PABF1 056** | ATAAGACACACACGCACATCTAG GGCGGCGGATTTGTCC | **AGACACACA CGCACAT** |
| | **PABR1 056** | AATATGTGCGTGTGTGTCTGCGG CAACCGAGCGTTCTG | |
| **SEVA_COLONY 2 / Sample7** | **PABF1 057** | ATAGACGAGCGTCTGAGAGCTA GGGCGGCGGATTTGTCC | **GACGAGCGT CTGAGAG** |
| | **PABR1 057** | AATCTCTCAGACGCTCGTCGCGG CAACCGAGCGTTCTG | |
| **SEVA_COLONY 3 / Sample8** | **PABF1 058** | ATATGTGTCTCTGAGAGTACTAG GGCGGCGGATTTGTCC | **TGTGTCTCTG AGAGTA** |
| | **PABR1 058** | AATTACTCTCAGAGACACAGCGG CAACCGAGCGTTCTG | |
| **SEVA_COLONY 4 / Sample9** | **PABF1 059** | ATACACACGCACTGAGATACTAG GGCGGCGGATTTGTCC | **CACACGCAC TGAGATA** |
| | **PABR1 059** | AATTATCTCAGTGCGTGTGGCGG CAACCGAGCGTTCTG | |
| **SEVA_COLONY 5 / Sample10** | **PABF1 060** | ATAGATGAGTATAGACACACTAG GGCGGCGGATTTGTCC | **GATGAGTAT AGACACA** |
| | **PABR1 060** | AATTGTGTCTATACTCATCGCGG CAACCGAGCGTTCTG | |
| **SEVA_COLONY 6 / Sample11** | **PABF1 061** | ATAGCTGTGTGTGCTCGTCCTAG GGCGGCGGATTTGTCC | **GCTGTGTGT GCTCGTC** |
| | **PABR1 061** | AATGACGAGCACACACAGCGCG GCAACCGAGCGTTCTG | |
| **SEVA_COLONY 7 / Sample12** | **PABF1 062** | ATATCTCAGATAGTCTATACTAG GGCGGCGGATTTGTCC | **TCTCAGATA GTCTATA** |
| | **PABR1 062** | AATTATAGACTATCTGAGAGCGG CAACCGAGCGTTCTG | |
| **SEVA_COLONY 8 / Sample13** | **PABF1 063** | ATAACACGCATGACACACTCTAG GGCGGCGGATTTGTCC | **ACACGCATG ACACACT** |
| | **PABR1** | AATAGTGTGTCATGCGTGTGCGG | |

| | | | |
|---|---|---|---|
| | 063 | CAACCGAGCGTTCTG | |
| **SEVA_COLONY 9 / Sample14** | **PABF1 064** | ATATATATACAGAGTCGAGCTAG GGCGGCGGATTTGTCC | **TATATACAG AGTCGAG** |
| | **PABR1 064** | AATCTCGACTCTGTATATAGCGG CAACCGAGCGTTCTG | |
| **SEVA_COLONY 10 / Sample15** | **PABF1 065** | ATAGCGCTCTCTCACATACCTAG GGCGGCGGATTTGTCC | **GCGCTCTCT CACATAC** |
| | **PABR1 065** | AATGTATGTGAGAGAGCGCGCG GCAACCGAGCGTTCTG | |

**Table S9: Results of the RBS strength calculation for the original (WT) and the mutated (pGreenL) RBS regions of *ccaS* and *ccaR* genes using Salis software*.**

| gene RBS | Translation rate | $\Delta G_{TOTAL}$ | $\Delta G_{mRNA\text{-}rRNA}$ | $\Delta G_{spacing}$ | $\Delta G_{stacking}$ | $\Delta G_{standby}$ | $\Delta G_{start}$ | $\Delta G_{mRNA}$ |
|---|---|---|---|---|---|---|---|---|
| ***ccaR* WT** | 7.18 | 11.44 | -13.30 | 1.52 | 0.00 | 2.56 | -0.42 | -20.96 |
| ***ccaR* pGreenL** | 20.30 | 9.13 | -14.64 | 1.52 | 0.00 | 2.56 | -0.42 | -19.99 |
| ***ccaS* WT** | 39.84 | 7.63 | -12.82 | 1.52 | 0.00 | 4.48 | -0.42 | -15.12 |
| ***ccaS* pGreenL** | 2.15 | 14.12 | -12.27 | 1.52 | 0.00 | 5.74 | -0.42 | -19.80 |

*https://salislab.net/software/predict_operon_calculator

### pGreenL sequence

ORFS are marked in bold, regulatory regions are underlined and mutations are marked in red

GGGCGCGCCGTAGAAAAGATCAAAGGATCTTCTTGAGATCCTTTTTTTCTGCGGGGGATCAGGACC
GCTGCCGGAGCGCAACCCACTCACTACAGCAGAGCCATGTAGGGCCGCCGGCGTTGTGGATACCTC
GCGGAAAACTTGGCCCTCACTGACAGATGAGGGGCGGACGTTGACACTTGAGGGGCCGACTCACC
CGGCGCGGCGTTGACAGATGAGGGGCAGGCTCGATTTCGGCCGGCGACGTGGAGCTGGCCAGCCT
CGCAAATCGGCGAAAACGCCTGATTTTACGCGAGTTTCCCACAGATGATGTGGACAAGCCTGGGGA
TAAGTGCCCTGCGGTATTGACACTTGAGGGGCGCGACTACTGACAGATGAGGGGCGCGATCCTTG
ACACTTGAGGGGCAGAGTGCTGACAGATGAGGGGCGCACCTATTGACATTTGAGGGGCTGTCCAC
AGGCAGAAAATCCAGCATTTGCAAGGGTTTCCGCCCGTTTTTCGGCCACCGCTAACCTGTCTTTTAA
CCTGCTTTTAAACCAATATTTATAAACCTTGTTTTTAACCAGGGCTGCGCCCTGTGCGCGTGACCGCG
CACGCCGAAGGGGGGTGCCCCCCCTTCTCGAACCCTCCCGGCCCGCTAACGCGGGCCTCCCATCCC
CCCAGGGGCTGCGCCCCTCGGCCGCGAACGGCCTCACCCCAAAAATGGCAGCCACGTAGAAAGCC
AGTCCGCAGAAACGGTGCTGACCCCGGATGAATGTCAGCTACTGGGCTATCTGGACAAGGGAAAA
CGCAAGCGCAAAGAGAAAGCAGGTAGCTTGCAGTGGGCTTACATGGCGATAGCTAGACTGGGCGG
TTTTATGGACAGCAAGCGAACCGGAATTGCCAGCTGGGGCGCCCTCTGGTAAGGTTGGGAAGCCCT
GCAAAGTAAACTGGATGGCTTTCTTGCCGCCAAGGATCTGATGGCGCAGGGGATCAAGATCGACG
GATCGATCCGGGGAATTAATTCCGGGGCAATCCCGCAAGGAGGGTGA**ATGAATCGGACGTTTGAC**
**CGGAAGGCATACAGGCAAGAACTGATCGACGCGGGGTTTTCCGCCGAGGATGCCGAAACCATCG**
**CAAGCCGCACCGTCATGCGTGCGCCCCGCGAAACCTTCCAGTCCGTCGGCTCGATAGTCCAGCAA**
**GCTACGGCCAAGATCGAGCGCGACAGCGTGCAACTGGCTCCCCCTGCCCTGCCCGCGCCATCGGC**
**CGCCGTGGAGCGTTCGCGTCGTCTCGAACAGGAGGCGGCAGGTTTGGCGAAGTCGATGACCATC**
**GACACGCGAGGAACTATGACGACCAAGAAGCGAAAAACCGCCGGCGAGGACCTGGCAAAACAG**
**GTCAGCGAAGCCAAGCAGGCCGCGTTGCTGAAACACACGAAGCAGCAGATCAAGGAAATGCAG**
**CTTTCCTTGTTCGATATTGCGCCGTGGCCGGACACGATGCGAGCGATGCCAAACGACACGGCCCG**
**CTCTGCCCTGTTCACCACGCGCAACAAGAAAATCCCGCGCGAGGCGCTGCAAAACAAGGTCATTT**
**TCCACGTCAACAAGGACGTGAAGATCACCTACACCGGCGTCGAGCTGCGGGCCGACGATGACGA**
**ACTGGTGTGGCAGCAGGTGTTGGAGTACGCGAAGCGCACCCCTATCGGCGAGCCGATCACCTTCA**
**CGTTCTACGAGCTTTGCCAGGACCTGGGCTGGTCGATCAATGGCCGGTATTACACGAAGGCCGAG**
**GAATGCCTGTCGCGCCTACAGGCGACGGCGATGGGCTTCACGTCCGACCGCGTTGGGCACCTGGA**
**ATCGGTGTCGCTGCTGCACCGCTTCCGCGTCCTGGACCGTGGCAAGAAAACGTCCCGTTGCCAGG**
**TCCTGATCGACGAGGAAATCGTCGTGCTGTTTGCTGGCGACCACTACACGAAATTCATATGGGAG**
**AAGTACCGCAAGCTGTCGCCGACGGCCCGACGGATGTTCGACTATTTCAGCTCGCACCGGGAGCC**
**GTACCCGCTCAAGCTGGAAACCTTCCGCCTCATGTGCGGATCGGATTCCACCCGCGTGAAGAAGT**
**GGCGCGAGCAGGTCGGCGAAGCCTGCGAAGAGTTGCGAGGCAGCGGCCTGGTGGAACACGCCT**
**GGGTCAATGATGACCTGGTGCATTGCAAACGCTAG**GGCCTTGTGGGGTCAGTTCCGGCTGGGGGT
TCAGCAGCCACCTGCATCGCGGCCGGCCTACGGCCAGCCTCGCAGAGCAGGATTCCCGTTGAGCAC
CGCCAGGTGCGAATAAGGGACAGTGAAGAAGGAACACCCGCTCGCGGGTGGGCCTACTTCACCTA
TCCTGCCCGGCTGACGCCGTTGGATACACCAAGGAAAGTCTACACGAACCCTTTGGCAAAATCCTGT
ATATCGTGCGAAAAAGGATGGATATACCGAAAAAATCGCTATAATGACCCCGAAGCAGGGTT**ATGC**
**AGCGGAAAAGGACAACGCGCGGACCGTTGTCCAATTTACCGAACAACTCCGCGGCCGGGAAGCC**
**GATCTCGGCTTGAACGAATTGTTAGGTGGCGGTACTTGGGTCGATATCAAAGTGCATCACTTCTTC**
**CCGTATGCCCAACTTTGTATAGAGAGCCACTGCGGGATCGTCACCGTAATCTGCTTGCACGTAGAT**
**CACATAAGCACCAAGCGCGTTGGCCTCATGCTTGAGGAGATTGATGAGCGCGGTGGCAATGCCCT**

**GCCTCCGGTGCTCTCCGGAGACTGCGAGATCATAGATATAGATCTCACTACGCGGCTGCTCAAACT**
**TGGGCAGAACGTAAGCCGCGAGAGCGCCAACAACCGCTTCTTGGTCGAAGGCAGCAAGCGCGAT**
**GAATGTCTTACTACGGAGCAAGTTCCCGAGGTAATCGGAGTCCGGCTGATGTTGGGAGTAGGTG**
**GCTACGTCTCCGAACTCACGACCGAAAAGATCAAGAGCAGCCCGCATGGATTTGACTTGGTCAGG**
**GCCGAGCCTACATGTGCGAATGATGCCCATACTTGAGCCACCTAACTTTGTTTTAGGGCGACTGCC**
**CTGCTGCGTAACATCGTTGCTGCTGCGTAACATCGTTGCTGCTCCATAACATCAAACATCGACCCA**
**CGGCGTAACGCGCTTGCTGCTTGGATGCCCGAGGCATAGGCTGTACAAAAAAACAGTCATAACAA**
**GCCATGAAAACCGCCACTGCGCCGTTACCACCGCTGCGTTCGGTCAAGGTTCTGGACCAGTTGCGT**
**GAGCGCATACGCTACTTGCATTACAGTTTACGAACCGAACAGGCTTATGTCAAATTTAA**ATCGTAA
TTATTGGGGACCCCTGGATTCTCACCAATAAAAAACGCCCGGCGGCAACCGAGCGTTCTGAACAAA
TCCAGATGGAGTTCTGAGGTCATTACTGGATCTATCAACAGGAGTCCAAGACTAGTCGCCAGGGTT
TTCCCAGTCACGACGCGGCCGCAAGCTTTCAAAAACCCCAGTTTTTACGGCTAGCTCAGTCCTAGGT
ATAGTGCTAGCTATAGAGGTTCAGAGCTTAGTACGAGAAGGCGTA**ATGGGCAAATTTCTAATTCCA**
**ATCGAATTTGTTTTTCTGGCGATCGCCATGACCTGTTATTTATGGCACAGACAAAACCAAGAACGC**
**CGCAGGATTGAAATTAGCATCAAGCAACAAACCCAACGGGAACGATTTATTAACCAAATTACCCA**
**ACATATCCGCCAATCTTTAAACTTGGAAACGGTTTTAAATACCACCGTCGCTGAAGTTAAAACCCT**
**GTTGCAAGTTGATCGAGTTCTAATTTATCGCATTTGGCAAGATGGCACGGGCAGCGCCATTACGG**
**AATCGGTGAATGCCAATTATCCTAGTATTTTAGGGCGGACCTTTTCCGATGAAGTTTTTCCCGTTG**
**AATACCATCAAGCCTACACCAAAGGTAAAGTACGGGCCATTAATGACATTGACCAGGATGACATA**
**GAGATTTGCCTAGCTGATTTCGTCAAACAATTTGGCGTGAAATCAAAATTAGTAGTGCCCATTCTT**
**CAACATAATCGTGCTTCTTCCCTAGATAATGAATCAGAATTTCCCTATCTTTGGGGGCTGTTAATTA**
**CCCATCAATGTGCTTTTACCCGGCCATGGCAACCGTGGGAAGTGGAGTTAATGAAACAGCTAGCC**
**AATCAGGTCGCGATCGCCATCCAACAATCGGAATTATATGAGCAATTACAGCAACTCAATAAAGA**
**TTTGGAAAACCGAGTCGAAAAACGCACCCAGCAACTTGCCGCCACCAATCAATCCCTAAGAATGG**
**AAATCAGTGAGCGACAAAAAACGGAAGCCGCTCTCCGCCACACTAACCATACTCTGCAATCCCTG**
**ATTGCGGCCTCCCCCAGGGGTATTTTTACCCTTAATTTAGCAGACCAAATTCAGATTTGGAATCCTA**
**CAGCAGAACGTATTTTTGGTTGGACAGAAACAGAAATTATTGCCCATCCAGAATTATTAACATCCA**
**ACATTTTGCTGGAAGATTATCAGCAATTTAAACAGAAAGTTTTATCAGGCATGGTTTCCCCTAGCC**
**TAGAATTAAAATGTCAAAAAAAAGATGGTAGTTGGATTGAAATTGTCCTTTCCGCTGCTCCCCTAT**
**TGGATAGTGAAGAAAATATTGCCGGATTGGTGGCGGTTGTCGCCGATATTACCGAGCAAAAGCG**
**GCAGGCAGAACAAATTCGTTTGCTACAATCCGTTGTGGTTAATACTAATGATGCGGTGGTGATTA**
**CGGAAGCGGAGCCCATTGATGATCCCGGGCCGAGAATTCTCTATGTCAATGAAGCATTTACTAAA**
**ATCACCGGTTATACTGCTGAAGAAATGCTAGGCAAAACCCCCCGAGTTTTACAGGGACCAAAAAC**
**TAGTCGCACTGAATTAGATAGGGTGCGGCAAGCCATTAGTCAATGGCAATCAGTTACCGTTGAAG**
**TGATTAATTATCGTAAGGATGGCAGTGAGTTTTGGGTGGAATTTAGTCTGGTGCCCGTTGCCAAT**
**AAAACAGGTTTTTACACCCATTGGATTGCTGTGCAAAGGGATGTCACTGAGCGCCGACGCACGGA**
**GGAAGTCCGCCTAGCTTTAGAACGGGAAAAAGAATTAAGCCGCCTAAAAACTCGTTTTTTCTCCA**
**TGGCTTCCCATGAATTTCGTACTCCCCTCAGTACGGCCTTAGCTGCTGCCCAATTACTGGAAAATTC**
**TGAAGTGGCCTGGCTTGATCCCGATAAGCGTAGCCGGAACTTACACCGTATTCAAAATTCCGTGA**
**AAAATATGGTACAGCTCCTGGATGATATTTTAATCATTAACCGTGCCGAAGCGGGCAAATTGGAA**
**TTTAATCCTAATTGGTTAGATTTGAAATTATTGTTCCAGCAATTTATCGAAGAAATTCAATTAAGT**
**GTCAGTGACCAATATTATTTTGACTTTATTTGTAGCGCTCAAGATACGAAGGCATTGGTGGATGA**
**AAGGTTAGTGCGGTCTATTTTATCTAATCTGTTATCTAATGCGATTAAATACTCTCCCGGGGGAGG**
**GCAGATTAAAATTGCCCTAAGCCTAGATTCGGAACAGATTATTTTTGAAGTCACCGACCAGGGCA**
**TTGGCATTTCGCCAGAGGACCAAAAGCAAATTTTTGAACCCTTTCATCGGGGCAAAAATGTCAGA**

**AATATTACGGGAACAGGACTCGGTTTAATGGTTGCCAAGAAATGTGTTGACTTACACAGTGGCAGTATCTTGCTAAAAAGTGCAGTTGACCAGGGAACAACAGTTACTATCTGTTTAAAACGCTATAACCATTTGCCTCGAGCTTAG**TAATAATCTAGACCAGGCATCAAATAAAACGAAAGGCTCAGTCGAAAGACTGGGCCTTTCGTTTTATCTGTTGTTTGTCGGTGAACGCTCTCTACTAGAGTCACACTGGCTCACCTTCGGGTGGGCCTTTCTGCGTTTATAGTAAATCACTGCATAATTCGTGTCGATTAAACCGCCTAAGTCCCCCAGGAAAGGGGGATATAACAGTATAGATTTTGTCAGCCTTCAGCTTGGCTTTACCGTCAAAAAAACTGACAGCTAGCTCAGTCCTAGGTATAATGCTAGCTCACACAGAATTCATTAAAGAGGAGAAAGGTACC**ATGAGTGTCAACTTAGCTTCCCAGTTGCGGGAAGGGACGAAAAAATCCCACTCCATGGCGGAGAACGTCGGCTTTGTCAAATGCTTCCTCAAGGGCGTTGTCGAGAAAAATTCCTACCGTAAGCTGGTTGGCAATCTCTACTTTGTCTACAGTGCCATGGAAGAGGAAATGGCAAAATTTAAGGACCATCCCATCCTCAGCCACATTTACTTCCCCGAACTCAACCGCAAACAAAGCCTAGAGCAAGACCTGCAATTCTATTACGGCTCCAACTGGCGGCAAGAAGTGAAAATTTCTGCCGCTGGCCAAGCCTATGTGGACCGAGTCCGGCAAGTGGCCGCTACGGCCCCTGAATTGTTGGTGGCCCATTCCTACACCCGTTACCTGGGGGATCTTTCCGGCGGTCAAATTCTCAAGAAAATTGCCCAAAATGCCATGAATCTCCACGATGGTGGCACAGCTTTCTATGAATTTGCCGACATTGATGACGAAAAGGCTTTTAAAAATACCTACCGTCAAGCTATGAATGATCTGCCCATTGACCAAGCCACCGCCGAACGGATTGTGGATGAAGCCAATGACGCCTTTGCCATGAACATGAAAATGTTCAACGAACTTGAAGGCAACCTGATCAAGGCGATCGGCATTATGGTGTTCAACAGCCTCACCCGTCGCCGCAGTCAAGGCAGCACCGAAGTTGGCCTCGCCACCTCCGAAGGCTAG**TTAAAGAGGAGAAAGGATCC**ATGGCCGTCACTGATTTAAGTTTGACCAATTCTTCCCTGATGCCCACGTTGAACCCGATGATTCAACAGTTGGCCCTGGCGATCGCCGCTAGTTGGCAAAGTTTACCCCTCAAGCCCTATCAATTGCCGGAGGATTTGGGCTACGTAGAAGGCCGCCTGGAAGGGGAAAAGTTAGTGATTGAAAATCGGTGCTACCAAACGCCCCAGTTTCGCAAAATGCATTTGGAGTTGGCCAAGGTGGGCAAAGGGTTGGATATTCTCCACTGTGTAATGTTTCCTGAGCCTTTATACGGTCTACCTTTGTTTGGCTGTGACATTGTGGCCGGCCCCGGTGGAGTAAGTGCGGCTATTGCGGATCTATCCCCCACCCAAAGCGATCGCCAATTGCCCGCAGCGTACCAAAAATCATTGGCAGAGCTAGGCCAGCCAGAATTTGAGCAACAACGGGAATTGCCCCCCTGGGGAGAAATATTTTCTGAATATTGTTTATTCATCCGTCCCAGCAATGTCACTGAAGAAGAAAGATTTGTACAAAGGGTAGTGGACTTTTTGCAAATTCATTGTCACCAATCCATCGTTGCCGAACCCTTGTCTGAAGCTCAAACTTTGGAGCACCGTCAGGGGCAAATTCATTACTGCCAACAACAACAGAAAAATGATAAAACCCGTCGGGTACTGGAAAAAGCTTTTGGGGAAGCTTGGGCGGAACGGTATATGAGCCAAGTCTTATTTGATGTTATCCAATAA**TCTAGACCAGGCATCAAATAAAACGAAAGGCTCAGTCGAAAGACTGGGCCTTTCGTTTTATCTGTTGTTTGTCGGTGAACGCTCTCTACTAGAGTCACACTGGCTCACCTTCGGGTGGGCCTTTCTGCGTTTATAGGTACCCGGGGATCCTCTAGAGTCGACCTGCAGGCATGCAAGCTTAGCAAGCGAACCGGAATTGCCAGCTGGGGCGCCCTCTGGTAAGGTTGGGAAGCCCTGCAAAGTAATTGACGGCTAGCTCAGTCCTAGGTACAGTGCTAGCTGTTAAACGCCCCTCGTAAGAACGCGGGAAAGCA**ATGAGAATTCTTTTAGTGGAGGATGATTTGCCGCTGGCGGAAACCCTTGCTGAAGCATTGAGTGACCAGCTTTACACCGTTGATATTGCCACCGACGCTTCCCTCGCCTGGGACTATGCCTCCCGACTGGAATATGACCTCGTTATTTTGGATGTGATGCTGCCGGAGTTGGACGGGATTACCCTCTGTCAAAAATGGCGATCGCACAGTTATTTAATGCCAATTTTGATGATGACAGCCAGGGATACGATCAATGATAAAATCACGGGCTTGGATGCGGGGGCGGATGATTATGTGGTCAAGCCAGTGGATTTGGGGGAGTTATTTGCCAGGGTGCGAGCTTTGTTGCGTCGGGGTTGTGCAACGTGCCAACCAGTTTTAGAGTGGGGGCCAATCAGGTTGGATCCAAGCACCTATGAAGTTAGTTATGACAATGAGGTTTTGTCTTTGACCCGCAAGGAATACAGCATTCTGGAATTACTACTCCGCAATGGCCGTCGGGTGCTAAGTCGGAGCATGATTATCGATAGTATCTGGAAGTTGGAGAGTCCCCCAGAGGAAGATACGGTTAAGGTGCATGTGCGGAGTTTGCGACAAAAATTAAAAAGTGCCGGTTTATCAGCAGATGCCATTGAAACGGTCCATGGCATTGGGTATCGTCTGGCCAATTTAACGG**

**AAAAATCTTTGTGCCAAGGGAAAAACTAG**TAATAATCTAGACCAGGCATCAAATAAAACGAAAGGCTCAGTCGAAAGACTGGGCCTTTCGTTTTATCTGTTGTTTGTCGGTGAACGCTCTCTACTAGAGTCACACTGGCTCACCTTCGGGTGGGCCTTTCTGCGTTTATAGTAAATCACTGCATAATTCGTGTCGCTCAAGGCGCACTCCCGTTCTGGATAATGTTTTTTGCGCCGACATCATAACGGTTCTGGCAAATATTCTGAAATGAGCTGGTTAGCTAGTCAAGCCCATTGTGCTTTTCTCTATCAACCTCAGCTTACCTGAAGGGGTGAACAGGTCTGGGTTAATTCATGTTGCGAAATGTAACAGTTTTAGTCGCATCAGCTAACTTTCCGATTTCTTTACGATTTTCTCCCCCTTTTCTTCAATTTTACTTTGTTAGGATCGCATTTTTAAAAAGAGGAGAAATACTAG**ATGCGTAAAGGCGAAGAGCTGTTCACTGGTGTCGTCCCTATTCTGGTGGAACTGGATGGTGATGTCAACGGTCATAAGTTTTCCGTGCGTGGCGAGGGTGAAGGTGACGCAACTAATGGTAAACTGACGCTGAAGTTCATCTGTACTACTGGTAAACTGCCGGTACCTTGGCCGACTCTGGTAACGACGCTGACTTATGGTGTTCAGTGCTTTGCTCGTTATCCGGACCATATGAAGCAGCATGACTTCTTCAAGTCCGCCATGCCGGAAGGCTATGTGCAGGAACGCACGATTTCCTTTAAGGATGACGGCACGTACAAAACGCGTGCGGAAGTGAAATTTGAAGGCGATACCCTGGTAAACCGCATTGAGCTGAAAGGCATTGACTTTAAAGAAGACGGCAATATCCTGGGCCATAAGCTGGAATACAATTTTAACAGCCACAATGTTTACATCACCGCCGATAAACAAAAAAATGGCATTAAAGCGAATTTTAAAATTCGCCACAACGTGGAGGATGGCAGCGTGCAGCTGGCTGATCACTACCAGCAAAACACTCCAATCGGTGATGGTCCTGTTCTGCTGCCAGACAATCACTATCTGAGCACGCAAAGCGTTCTGTCTAAAGATCCGAACGAGAAACGCGATCATATGGTTCTGCTGGAGTTCGTAACCGCAGCGGGCATCACGCATGGTAT**GGATGAACTGTACAAATGATGATTAAAGGCATCAAATAAAACGAAAGGCTCAGTCGAAAGACTGGGCCTTTCGTTTTATCTGTTGTTTGTCGGTGAACGCTCTCCTGAGTAGGACAAATCCGCCGCCCTAGACAGCT
